# SARS-CoV-2 infection of olfactory epithelial cells and neurons drives acute lung injury and lethal COVID-19 in mice

**DOI:** 10.1101/2021.12.04.471245

**Authors:** Alan T. Tang, David W. Buchholz, Katherine M. Szigety, Brian Imbhiaka, Siqi Gao, Maxwell Frankfurter, Min Wang, Jisheng Yang, Peter Hewins, Patricia Mericko-Ishizuka, N Adrian Leu, Stephanie Sterling, Isaac A. Monreal, Julie Sahler, Avery August, Xuming Zhu, Kellie A. Jurado, Mingang Xu, Edward E. Morrisey, Sarah E. Millar, Hector C. Aguilar, Mark L. Kahn

## Abstract

Lethal COVID-19 is associated with respiratory failure that is thought to be caused by acute respiratory distress syndrome (ARDS) secondary to pulmonary infection. To date, the cellular pathogenesis has been inferred from studies describing the expression of ACE2, a transmembrane protein required for SARS-CoV-2 infection, and detection of viral RNA or protein in infected humans, model animals, and cultured cells. To functionally test the cellular mechanisms of COVID-19, we generated *hACE2*^fl^ animals in which human ACE2 (hACE2) is expressed from the mouse *Ace2* locus in a manner that permits cell-specific, Cre-mediated loss of function. *hACE2*^fl^ animals developed lethal weight loss and hypoxemia within 7 days of exposure to SARS-CoV-2 that was associated with pulmonary infiltrates, intravascular thrombosis and patchy viral infection of lung epithelial cells. Deletion of hACE2 in lung epithelial cells prevented viral infection of the lung, but not weight loss, hypoxemia or death. Inhalation of SARS-CoV-2 by *hACE2*^fl^ animals resulted in early infection of sustentacular cells with subsequent infection of neurons in the neighboring olfactory bulb and cerebral cortex— events that did not require lung epithelial cell infection. Pharmacologic ablation of the olfactory epithelium or *Foxg1*^Cre^ mediated deletion of hACE2 in olfactory epithelial cells and neurons prevented lethality and neuronal infection following SARS-CoV-2 infection. Conversely, transgenic expression of hACE2 specifically in olfactory epithelial cells and neurons in *Foxg1*^Cre^; LSL-*hACE2* mice was sufficient to confer neuronal infection associated with respiratory failure and death. These studies establish mouse loss and gain of function genetic models with which to genetically dissect viral-host interactions and demonstrate that lethal disease due to respiratory failure may arise from extrapulmonary infection of the olfactory epithelium and brain. Future therapeutic efforts focused on preventing olfactory epithelial infection may be an effective means of protecting against severe COVID-19.

## Introduction

Understanding the cellular mechanisms that underlie coronavirus disease 2019 (COVID-19) is necessary to determine how to best prevent infection and treat affected individuals. To date, COVID-19 pathogenesis has been inferred from descriptive studies, in particular those that identify the cell types that express angiotensin-converting enzyme 2 (ACE2), a transmembrane protein host receptor required for infection with the causative viral pathogen: Severe Acute Respiratory Syndrome Coronavirus 2 (SARS-CoV-2)^1, 2^, or viral RNA or protein in the tissues of infected individuals^3^ or animals^4–7^. These studies have supported diverse disease mechanisms that include both direct cell infection by the SARS-CoV-2 virus and indirect effects of systemic inflammatory signals. Defining causal mechanisms for COVID-19 using functional in vivo approaches would better identify cellular targets and inform rational design of optimized therapies and preventative strategies.

Common symptoms of acute COVID-19 include anosmia and, in the case of more severe disease, respiratory distress. Anosmia is thought to result from SARS-CoV-2 infection of olfactory epithelium (OE)^8, 9^, which is not believed to be a central mechanism of lethality associated with COVID-19 disease^10, 11^. In contrast, hypoxemic respiratory failure is the most common cause of death^12^. COVID-19 pneumonia has been proposed to arise through direct infection of lung epithelial cells and/or lung macrophages with progression to acute respiratory distress syndrome (ARDS), a non-cardiogenic pulmonary edema that may arise in association with systemic inflammatory signals (e.g. from pancreatitis) rather than a primary pulmonary insult^13, 14^. Testing and defining the roles of these distinct, proposed mechanisms is essential for effective prevention and treatment.

The application of mouse models to investigate COVID-19 pathogenesis has been hindered by the fact that that the SARS-CoV-2 spike protein is unable to bind the mouse ACE2 protein, a necessary first step in viral cellular entry and infection^1, 12^. Several human ACE2 (hACE2)-expressing mouse models have been generated to investigate COVID-19 pathogenesis^4, 7, 15, 16^, but only the K18-hACE2 line confers severe illness like that observed in patients^6^. K18-hACE2 random transgenic mice express hACE2 in a non-endogenous fashion that is primarily restricted to epithelial cells, and do not enable genetic dissection of viral pathogenesis. To identify cell specific roles during SARS-CoV-2 infection, we generated new lines of mice in which hACE2 is expressed from the mouse *Ace2* locus in a manner that enables cell-specific loss of function and mice in which hACE2 expression and gain of function can be conferred in a cell specific manner. Our results demonstrate that acute lung injury can occur in the absence of pulmonary infection and identify the olfactory epithelium and cerebral neurons as critical cellular sites of infection during lethal COVID-19.

## Results

### hACE2*^fl^* mice express the human ACE2 protein in an endogenous manner

To genetically investigate the cellular pathogenesis of COVID-19 disease *in vivo*, we generated gene targeted mice in which the coding sequence of the first translated exon of the mouse *Ace2* gene (exon 2) was replaced by an expression cassette including the human *ACE2* cDNA followed by a woodchuck hepatitis virus post-transcriptional regulatory element (WPRE) and simian-virus 40 (SV40) late polyA sequence (Fig. 1a). Since the start codon in Exon 2 is preceded by inter-species conserved 5’ untranslated sequence, we inserted loxP sites flanking the WPRE-polyA sequence to enable Cre-mediated loss of ACE2 expression without altering endogenous 5’ regulation of the *Ace2* locus. To compare hACE2 protein expression from the gene targeted allele (hereafter termed “*hACE2*^fl^”) and mouse ACE2 (mACE2) protein expression from the endogenous *Ace2* gene, we performed western blotting of whole tissue lysates from tissues known to express ACE2. Male mice were used because the *Ace2* allele is located on the X chromosome, enabling a straightforward comparison from the single expressed allele in all cell types of *hACE2*^fl/y^ animals. Western blotting with pan-ACE2 antibodies that recognize both hACE2 and mACE2 proteins revealed similar levels of ACE2 expression in heart, intestine, kidney and lung (Fig. 1b). To test the ability of Cre-mediated loss of the floxed WPRE-polyA cassette to block hACE2 protein expression, we generated germline, Cre-recombined animals (hACE2^del/y^) through intercrosses with Sox2-Cre animals. Western blotting with human ACE2-specific antibodies comparing wildtype, hACE2^fl/y^, and hACE2^del/y^ animals demonstrated loss of hACE2 protein in whole kidney and intestine (ileum) lysates (Fig. 1c).

**Figure 1.**
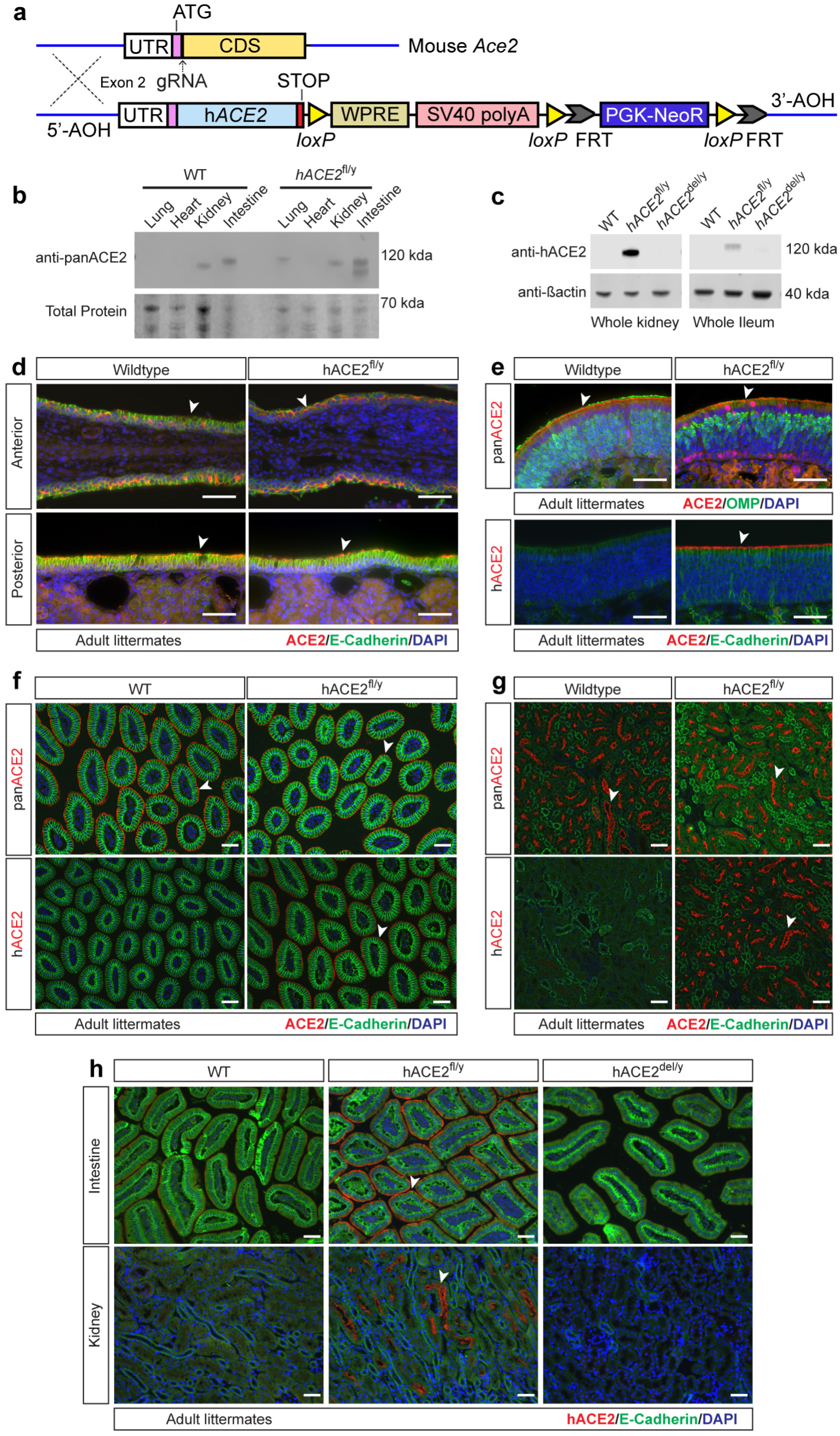
*hACE2*^fl/y^ knockin mice express human ACE2 in a conditional manner. **a**, Generation of the *hACE2^fl/y^* allele using gene targeting of the mouse *Ace2* locus. WPRE, woodchuck hepatitis virus post-transcriptional regulatory element. SV40, simian virus 40. PGK-NeoR, phosphoglycerate kinase promoter driven neomycin resistance cassette **b**, Immunoblotting of whole tissue lysates from wild-type (WT) and *hACE2*^fl/y^ mouse tissues was performed using an antibody recognizing both human and mouse ACE2 (panACE2). Total protein for each lane was detected using Revert counterstain. Representative of 3 independent experiments, n = 3 animals per genotype. **c**, Immunoblotting of whole tissue lysates from wild type (WT), *hACE2*^fl/y^ and *hACE2*^del/y^ tissues was performed using antibodies that specifically recognize human ACE2 (hACE2) protein. β-actin immunoblotting is shown as a loading control. Representative of 3 independent experiments, n = 3 animals per genotype. **d**, Immunohistochemistry using pan-ACE2 and E-Cadherin antibodies is shown for wild-type and *hACE2*^fl/y^ epithelium from the anterior and posterior part of the nasal cavity. Arrowheads indicate ACE2+ epithelial cells. n = 3 animals per genotype. **e**, Immunohistochemistry using pan-ACE2 and human-ACE2 (hACE2) antibodies is shown for wild-type and *hACE2*^fl/y^ olfactory epithelium co-stained with olfactory marker protein (OMP, mature olfactory sensory neurons) or epithelial cadherin (E-cadherin, sustentacular cells). Arrowheads indicate ACE2+ olfactory epithelial cells. n = 3 animals per genotype. **f**, Immunostaining using pan-ACE2 and human-ACE2 (hACE2) antibodies is shown for wild-type and *hACE2*^fl/y^ small intestine co-stained with E-cadherin (enterocytes). Arrowheads indicate ACE2+ enterocytes. n = 3 animals per genotype. **g**, Immunostaining using pan-ACE2 and human-ACE2 (hACE2) antibodies is shown for wild-type and *hACE2*^fl/y^ kidney co-stained with E-cadherin (epithelium). Arrowheads indicate ACE2+ tubular epithelial cells. n = 3 animals per genotype. Scale bars for all images, 50 µm.

To further compare expression and localization of hACE2 in knock-in animals and endogenous mACE2 expression, we next performed immunostaining of tissue sections with pan-ACE2 antibodies and antibodies specific for hACE2. ACE2 expression was detected in a discontinuous pattern on the apical brush border of ciliated epithelial cells in the anterior and posterior nasal respiratory epithelium of both *hACE2^fl^* and wild-type mice (Fig. 1d), a finding consistent with prior reports of its localization in ciliated epithelial cells^9, 17^. Immunostaining of epithelial cells in the olfactory epithelium, intestine and kidney using pan-ACE2 and human-specific ACE2 antibodies also showed similar expression levels and an apical cellular expression pattern in control and *hACE2^fl^* animals (Fig. 1e-g), while staining of the lung failed to reveal detectable ACE2 expression in either wild-type or *hACE2^fl^* mice (not shown). Lastly, immunostaining of *hACE2*^del/y^ tissues demonstrated loss of hACE2 expression in the intestine and kidney compared to littermate *hACE2*^fl/y^ tissues (Fig. 1h). These studies suggested that the *hACE2*^fl^ allele expresses hACE2 in a manner like that of the endogenous mouse *Ace2* allele, and that hACE2 protein expression is lost following Cre-mediated removal of the WPRE-polyA cassette.

### hACE2^fl^ mice experience lethal infection following exposure to SARS-CoV-2

A subset of human patients exposed to SARS-CoV-2 virus experience severe illness that leads to rapid respiratory failure and death^18, 19^. This severe disease course has been reproduced in K18-hACE2 transgenic mice that over-express hACE2 in epithelial cells throughout the respiratory tract^6^, but not in other animal models such as hamsters or previously reported hACE2 knockin mice^5, 7, 15, 20, 21^. To define the disease response of *hACE2*^fl^ mice, we exposed *hACE2*^fl/y^ and *hACE2*^del/y^ littermates to 10^5^ PFU of SARS-CoV-2 virus via nasal inhalation (Fig. 2a). *hACE2*^fl/y^ animals exhibited weight loss beginning 2 days after viral exposure and appeared lethargic, hunched and with ruffled fur by day 4, with sacrifice or death of all animals by 6 days post-infection due to >20% weight loss (humane endpoint) and/or respiratory distress (marked by tachypnea and flank heaving). In contrast, infected *hACE2*^del/y^ mice exhibited no weight loss or respiratory distress at timepoints up to 14 days post-infection (Fig. 2b, c), functionally confirming the loss of hACE2 function with Cre-recombination. Similar findings were obtained following exposure to 10^4^ PFU of SARS-CoV-2 virus (Fig. 2d, e), and following infection of *hACE2*^fl/y^ animals in two distinct ABSL3 facilities (Ext. Data Fig. 1). In contrast to *hACE2*^fl/y^ animals, a previously reported strain of hACE2 conditional knock-in animals^20^ failed to exhibit significant weight loss or lethality after exposure to 10^5^ SARS-CoV-2 virus (Fig. 2f, g).

**Figure 2.**
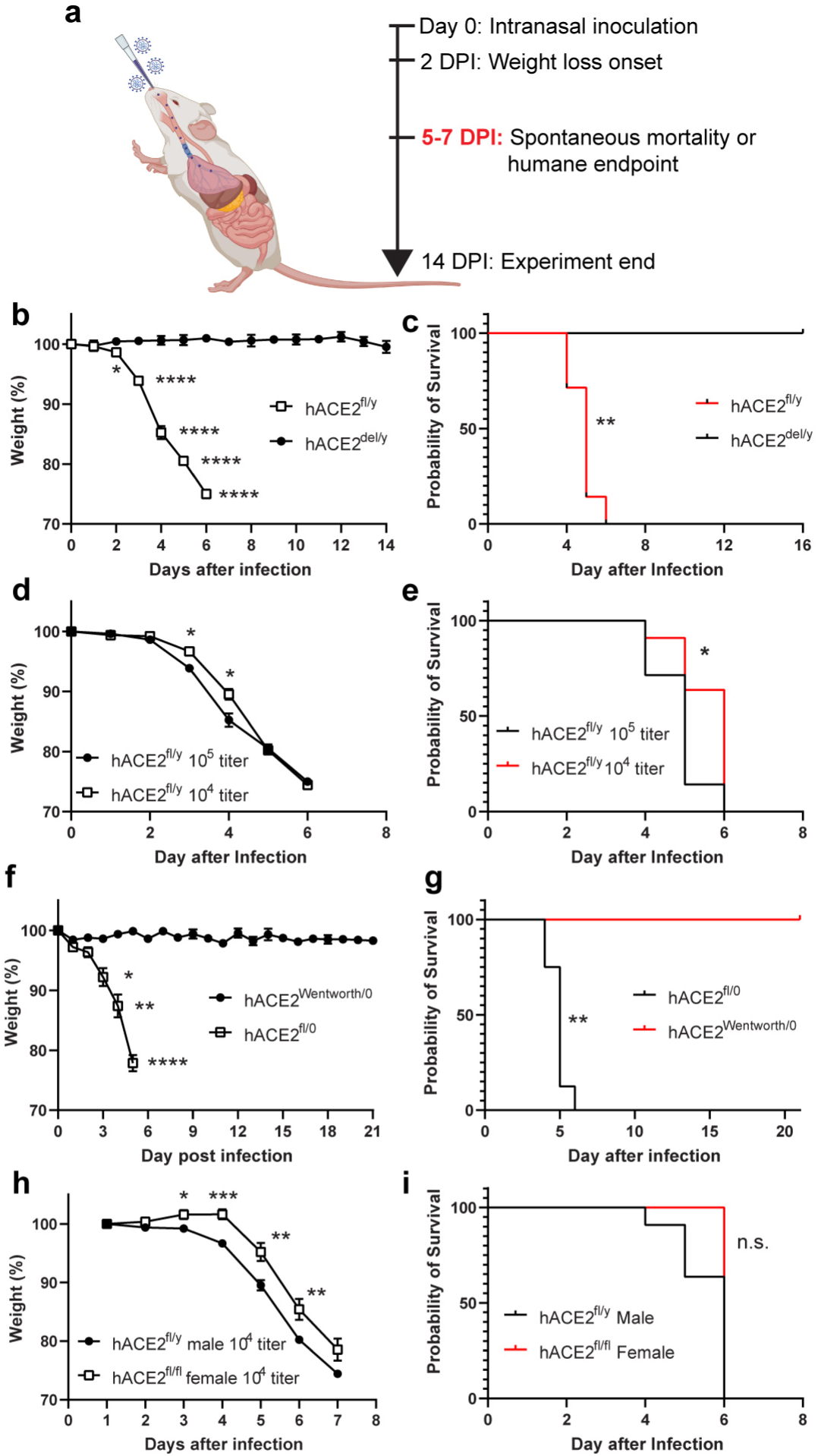
Intranasal SARS-CoV-2 infection of *hACE2*^fl/y^ mice confers weight loss and lethality reversed by Cre recombination. **a**, Experimental design for acute infection with SARS-CoV-2 virus. DPI, days post infection. **b-c**, Weight loss and survival of *hACE2*^fl/y^ and *hACE2*^del/y^ mice after infection with 10^5^ viral titer per mouse. n=8 for both genotypes, two independent experiments. **d-e**, Weight loss and survival of hACE2^fl/y^ mice after 10^5^ and 10^4^ titer infection of SARS-CoV-2. n=8 (10^5^ titer) and 11 (10^4^ titer), three independent experiments. **f-g**, Weight loss and survival of *hACE2*^fl/y^ and *hACE2*^Wentworth/y^ mice after infection with 10^5^ PFU of SARS-CoV-2 virus. n=8 (*hACE2*^fl/y^) and 5 (*hACE2*^Wentworth/y^), one experiment. **h-i**, Weight loss and survival of littermate male and female *hACE2^fl/y^* mice after infection with 10^4^ titer of SARS-CoV-2 virus. n= 11 (male) and 6 (female), one experiment. n.s p>0.05, *p<0.5, **p<0.01, ***p<0.001, ****p<0.0001 by multiple unpaired two-tailed t-test or log rank Mantel Cox test. Note: data for *hACE2*^fl/y^ infections at 10^5^ and 10^4^ titer are re-used throughout panels in this figure.

Human studies have revealed a sex difference in COVID-19 susceptibility, with male sex identified as an independent risk factor for severe illness and death^22, 23^. Consistent with these human findings, although severe weight loss and 100% mortality were observed in both groups male animals exhibited symptoms at earlier timepoints than females (Fig. 2h, i). Collectively, these studies demonstrate that *hACE2*^fl/y^ animals develop reproducible, severe disease and lethality after exposure to the SARS-CoV-2 virus.

### hACE2*^fl^* animals develop SARS-CoV-2 lung infection that requires epithelial cell hACE2 expression

Following infection with SARS-CoV-2, *hACE2*^fl/y^ animals exhibited labored breathing, suggestive of respiratory distress and COVID-19 lung disease. Prior studies using single cell RNAseq and immunohistochemical analysis of mouse and human lungs have identified surfactant protein C (SFTPC)-positive alveolar Type 2 (AT2) cells as ACE2-positive and suggested that AT2 cell infection might play a key role in COVID-19 pneumonia^24–27^. Therefore, we next used the SftpcCreERT2 knock-in allele to test whether ACE2-dependent infection of AT2 cells is required for lung infection in *hACE2*^fl^ animals. Following tamoxifen induction to activate Cre recombination, SftpcCreERT2; *hACE2*^fl/y^ males infected with SARS-CoV-2 virus exhibited levels of viral nucleocapsid protein and viral RNA in the lung that were indistinguishable from *hACE2*^fl/y^ animals by IHC and qPCR (Fig. 3a, b). These findings suggested that cell-types other than AT2 cells support SARS-CoV-2 infection of the lung, with the caveat that failure to detect hACE2 expression in AT2 cells precludes confirmation of its loss in those cells in SftpcCreERT2; *hACE2*^fl/y^ animals.

**Figure 3.**
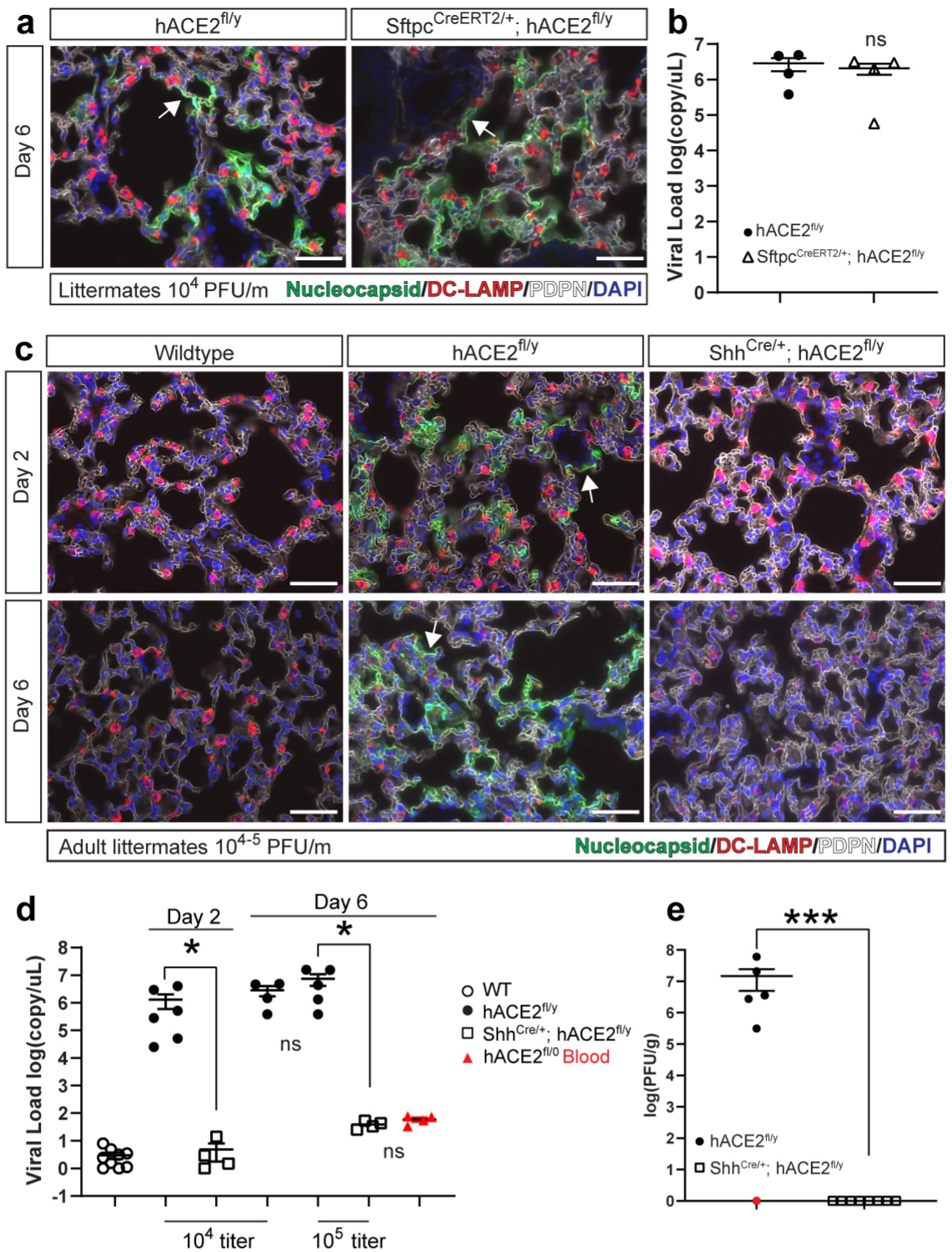
SARS-CoV-2 infects lung alveolar type I cells in an ACE2-dependent manner. **a**, Immunohistochemistry of *hACE2*^fl/y^ and Sfptc^CreERT2/+^; *hACE2*^fl/y^ mouse lungs 6 days after infection with 10^4^ PFU of SARS-CoV-2 virus was performed using antibodies that recognize SARS-CoV-2 nucleocapsid, DC-LAMP (alveolar type 2 cells) and PDPN (alveolar type 1 cells) as well as the nuclear stain DAPI. N=4 for both genotypes, one experiment **b**, qPCR was performed on *hACE2*^fl/y^ and Sfptc^CreERT2/+^; *hACE2*^fl/y^ mouse lungs harvested 6 days after infection with 10^4^ PFU of SARS-CoV-2 to measure total viral load. N=4 for both genotypes, one experiment. **c**, Immunohistochemistry of *hACE2*^fl/y^ and Shh^Cre/+^; *hACE2*^fl/y^ mouse lungs 2 and 6 days after infection with 10^4^ or 10^5^ PFU of SARS-CoV-2 virus was performed using antibodies that recognize SARS-CoV-2 nucleocapsid, DC-LAMP (alveolar type 2 cells) and PDPN (alveolar type 1 cells) as well as the nuclear stain DAPI. N=6 for all genotypes, two independent experiments. **d**, qPCR was performed on *hACE2*^fl/y^ and Shh^Cre/+^; *hACE2*^fl/y^ mouse lungs harvested 2 and 6 days after infection with 10^4^ or 10^5^ PFU of SARS-CoV-2 to measure total viral RNA load. Simultaneous measurement using whole blood was performed 6 days after infection (shown in red) to measure circulating levels. **e**, Infectious viral load was measured by plaque forming assay (PFU) from *hACE2*^fl/y^ and Shh^Cre/+^; *hACE2*^fl/y^ mouse lungs 6 days after infection with 10^5^ PFU of SARS-CoV-2. Scale bars, 50 µm. ***p<0.001; *p<0.05; ns p>0.05, significance determined by unpaired two-tailed t-test.

Histologic examination of *hACE2*^fl/y^ lungs 2 and 6 days after infection with SARS-CoV-2 revealed viral nucleocapsid staining primarily along the cell membrane of Podoplanin (PDPN)-positive alveolar Type 1 (AT1) epithelial cells (Fig. 3c). To test the role of ACE2 in both AT1 and AT2 cell infection, we examined SARS-CoV-2 infection of Shh^Cre^; *hACE2*^fl/y^ animals in which Cre is active in all lower respiratory and gastrointestinal epithelium (^28^ and Extended Data Figs. 2 and 3). SARS-CoV-2 infection was blocked completely in Shh^Cre^; *hACE2*^fl/y^ lungs, as assessed by immunostaining for viral nucleocapsid (Fig. 3c), measurement of lung viral load using both qPCR for viral genomic DNA (Fig. 3d) and culture of viral plaque forming units (PFUs) from lung tissue (Fig. 3e). A low level of virus was detected using qPCR on day 6 post-infection equivalent to the ambient level detected in circulating blood of *hACE2*^fl/y^ animals (Fig. 3d). These findings demonstrate that lung infection by SARS-CoV-2 occurs in both AT1 and AT2 epithelial cells in an ACE2-dependent manner.

### Lethal COVID-19 lung disease can arise without primary lung epithelial infection

Studies of human patients with severe COVID-19 pneumonia have revealed evidence of both lung infection by SARS-CoV-2 virus^3, 24, 25, 29^ and acute respiratory distress syndrome (ARDS), manifest by non-cardiogenic pulmonary edema, severe hypoxemia and a systemic inflammatory response^13, 30^. The extent to which these two mechanisms of lung disease are inter-connected and their respective roles in COVID-19 mortality have been difficult to define. Following exposure to 10^4^ PFU per mouse of SARS-CoV-2, both *hACE2*^fl/y^ and Shh^Cre^; *hACE2*^fl/y^ animals exhibited weight loss and lethality (Fig. 4a, b). There was a small delay in the onset of symptoms in Shh^Cre^; *hACE2*^fl/y^ animals, but both *hACE2*^fl/y^ and Shh^Cre^; *hACE2*^fl/y^ animals were severely hypoxic 5-6 days after infection, with oxygen (O2) saturations ranging between 70 and 85%, levels similar to those of severely ill patients requiring mechanical ventilation (Fig. 4c). H&E staining of lung sections from wildtype, *hACE2*^fl/y^ and Shh^Cre^*; hACE2*^fl/y^ animals 6 days after exposure to SARS-CoV-2 revealed alveolar consolidation, interstitial thickening and the presence of focal infiltrates and hemorrhage in both *hACE2*^fl/y^ and Shh^Cre^; *hACE2*^fl/y^ animals that were not observed in wildtype controls (Fig. 4d). Virtually identical results were obtained following exposure to 10^5^ PFU of SARS-CoV-2 (Ext. Data Fig. 4). SARS-CoV-2 infected *hACE2*^fl/y^ and Shh^Cre^; *hACE2*^fl/y^ animals also exhibited thrombus-filled pulmonary vessels that were not observed in the lungs of wild-type animals exposed to virus (Fig. 4d, e and Ext. Data Fig. 4), findings similar to those in human lungs harvested from individuals with lethal COVID-19 disease^13^.

**Figure 4.**
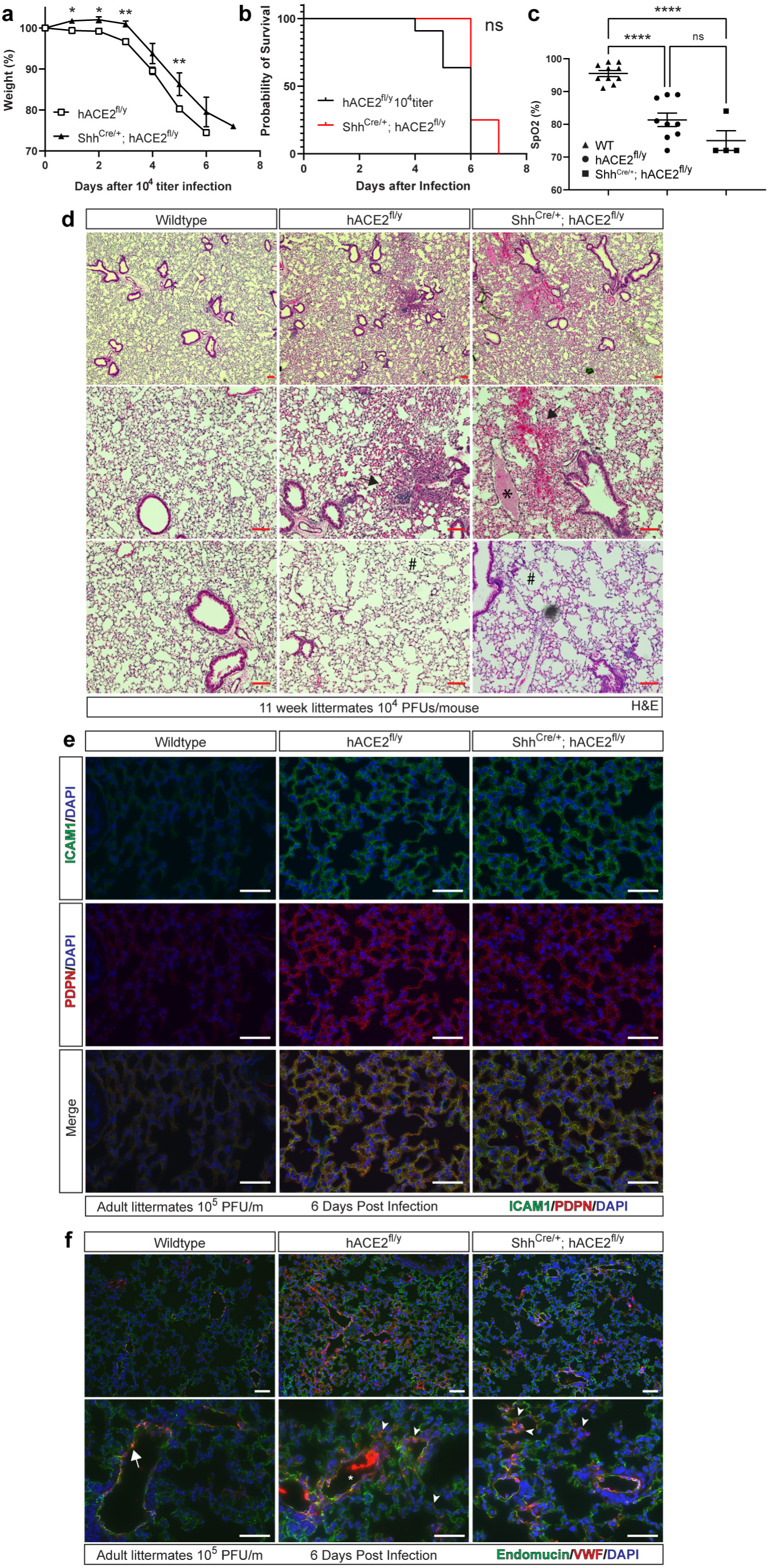
hACE2 mice develop acute lung injury and hypoxemia in the absence of SARS-CoV-2 lung infection. **a-b**, Weight loss and survival of *hACE2*^fl/y^ and Shh^Cre/+^; *hACE2*^fl/y^ mice after infection with 10^4^ PFU of SARS-CoV-2. N=11 (*hACE2*^fl/y^) and 4 (Shh^Cre/+^; *hACE2*^fl/y^), two independent experiments. Note: data for *hACE2*^fl/y^ same as Figure 2 since Shh^Cre/+^; *hACE2*^fl/y^ animals were littermates. **c**, Pulse oximetry measured in wild-type (WT), *hACE2*^fl/y^ and Shh^Cre/+^; *hACE2*^fl/y^ mice 6 days after exposure to 10^4^ PFU of SARS-CoV-2 virus. **d**, Hematoxylin-eosin (H&E) staining of wild-type (WT), *hACE2*^fl/y^ and Shh^Cre/+^; *hACE2*^fl/y^ lung tissue 6 days after exposure to 10^4^ PFU of SARS-CoV-2 virus. Arrows, sites of focal consolidation. Asterisks, intravascular thrombi. Hashtag, acute emphysematous changes. Representative of N=4 animals per genotype. **e-f**, Immunohistochemistry of wild-type (WT), *hACE2*^fl/y^ and Shh^Cre/+^; *hACE2*^fl/y^ lung tissue 6 days after exposure to 10^4^ PFU of SARS-CoV-2 virus using antibodies against Intracellular adhesion marker-1 (ICAM-1) and Podoplanin (PDPN), or von Willebrand’s Factor (vWF) and Endomucin (endothelial cells). Arrowheads in f identify vWF-positive microvasculature of the lung in *hACE2*^fl/y^ and Shh^Cre/+^; *hACE2*^fl/y^ animals. Representative of N=4 animals per genotype. Scale bars, 50 µm.

Consistent with a global lung inflammatory state, we observed uniformly elevated expression of the inflammation-induced proteins ICAM1 and Podoplanin (PDPN aka RTI40) in the alveolar epithelial cells of infected *hACE2*^fl/y^ and Shh^Cre^; *hACE2*^fl/y^ animals compared with SARS-CoV-2 inoculated wild-type controls (Fig. 4e)^31^. Expression of the pro-coagulant, inflammation-induced protein von Willebrand’s Factor (vWF) was also up-regulated in the lung capillary endothelial cells of SARS-CoV-2 infected *hACE2*^fl/y^ and Shh^Cre^; *hACE2*^fl/y^ animals compared with wild-type controls (Fig. 4f), consistent with the presence of intravascular thrombi in those lungs. These studies and those shown in Fig. 3 demonstrate that acute lung injury and hypoxemic respiratory failure may arise in the absence of primary SARS-CoV-2 lung infection, and implicate an inflammatory state and ARDS as the primary cause of respiratory failure and lethality in *hACE2*^fl/y^ mice.

### Lethal COVID-19 disease is associated with SARS-CoV-2 infection of the olfactory epithelium, olfactory bulb and cerebrum

The above results suggested that SARS-CoV-2 infection at cellular sites outside the lung and lacking Shh^Cre^ activity conferred severe disease in *hACE2*^fl/y^ animals. To define the extra-pulmonary sites of infection in Shh^Cre^; *hACE2*^fl/y^ mice that might be responsible for pulmonary inflammation and respiratory failure, we performed lineage tracing of the Shh^Cre^ transgene along the path of inhaled virus using the R26-LSL-RFP (Ai14) Cre reporter allele. Shh^Cre^ activity was detected uniformly in epithelial cells lining the trachea, lung airways and alveoli, and in isolated pockets of transition zone epithelium that lies between the nasal cavity respiratory and olfactory epithelium (Ext. Data Fig. 2b). In contrast, the respiratory epithelium (RE) and olfactory epithelium (OE) of the nasal passages failed to display Shh^Cre^ activity (Ext. Data Fig. 2b), suggesting that these sites of high ACE2 expression may be responsible for the lethal response to SARS-CoV-2 infection.

Therefore, we next compared SARS-CoV-2 nucleocapsid staining in the OE and adjacent olfactory bulb (OB) of the brain during the course of infection in wild-type, *hACE2*^fl/y^ and Shh^Cre^; *hACE2*^fl/y^ animals. 2 days after infection, abundant SARS-CoV-2 nucleocapsid was observed in the nasal respiratory and olfactory epithelium of *hACE2*^fl/y^ and Shh^Cre^; *hACE2*^fl/y^ animals, but none was detected in the adjacent OB and cerebral cortex of the brain (Fig. 5a, c). In contrast, 5-6 days after infection relatively little SARS-CoV-2 nucleocapsid was detected in the RE and OE of *hACE2*^fl/y^ and Shh^Cre^; *hACE2*^fl/y^ animals, although abundant signal was then detected in the neighboring OB, cerebral cortex, and hippocampus (Fig. 5b, d and Ext. Data Fig. 5). Co-staining with the nuclear neuronal marker NeuN and the glial cell marker GFAP revealed SARS-CoV-2 nucleocapsid specifically in neurons and not associated glial cells, and demonstrated the presence of a reactive gliosis 5 days post-infection (Fig. 5d). Sparse neuronal cell SARS-CoV-2 nucleocapsid and a reactive gliosis was also detected in the brainstem, a site more distant from the OE, but not in the cerebellum, 5 days after infection (Ext. Data Fig. 5). Consistent with the immunostaining studies to detect virus described above, in situ hybridization of the nasal area and adjacent tissues using RNAscope probes for SARS-CoV-2 RNA revealed little or no virus in the OE 5 days after infection, although abundant viral RNA was detected in the adjacent olfactory bulb (OB) and cerebrum of the brain at that timepoint (Fig. 5e, f and Ext. Data Fig. 6). These findings are consistent with prior studies demonstrating that olfactory epithelial cells may rapidly clear the virus after infection with SARS-CoV-2 in hamsters^32^ and in humans^9^, and suggest that early infection of the OE is followed by later infection of neurons in both neighboring and distal sites in the brain following SARS-CoV-2 exposure.

**Figure 5.**
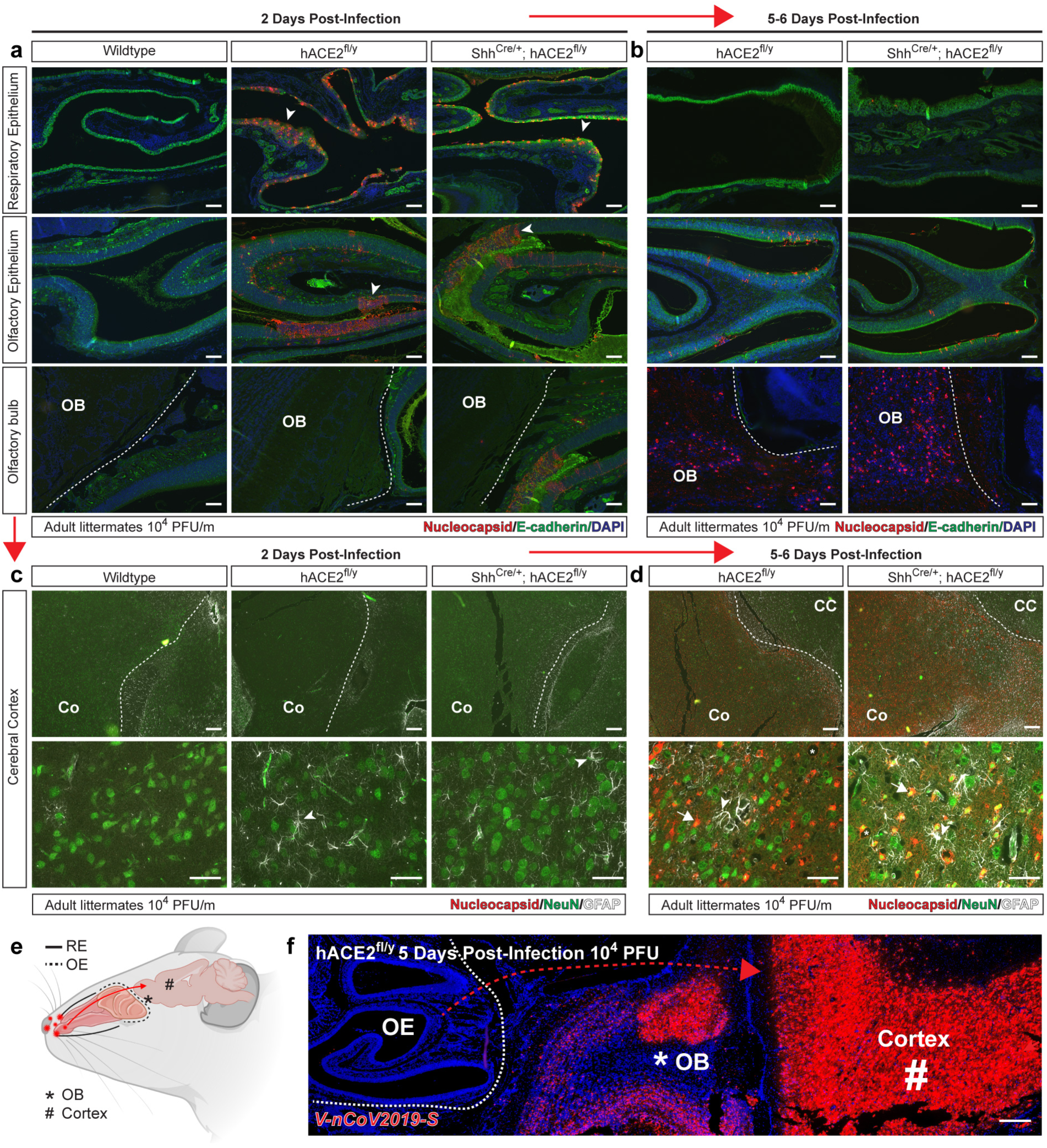
SARS-CoV-2 infection of the olfactory epithelium and brain in hACE2 mice. **a-b**, Immunohistochemistry of SARS-CoV-2 nucleocapsid and epithelial cell E-cadherin in the respiratory epithelium, olfactory epithelium, and olfactory bulb 2 and 5-6 days after infection of *hACE2*^fl/y^ and Shh^Cre/+^; *hACE2*^fl/y^ mice. Arrowheads indicate sites of viral nucleocapsid detection. Representative of N=4 animals per genotype and timepoint. Scale bars 100 µm. **c-d**, Immunohistochemistry of SARS-CoV-2 nucleocapsid, neuronal NeuN and glial cell GFAP in the cerebral cortex (Co) 2 and 5-6 days after infection. Arrowheads indicate sites of GFAP+ reactive gliosis. Arrows indicate nucleocapsid colocalization with NeuN staining. Representative of N=4 animals per genotype and timepoint. Scale bars 100 µm top, 50µm bottom. **e**, Diagram of the mouse nasal cavity and cranial anatomy. RE, respiratory epithelium; OE, olfactory epithelium; OB, olfactory bulb of the brain. **f**, *In situ* hybridization detection of SARS-CoV-2 mRNA 5 days post-infection reveals virus in the OB and cerebral cortex of the brain, but not the OE of the nose. Scale bar 250 µm.

**Figure 6.**
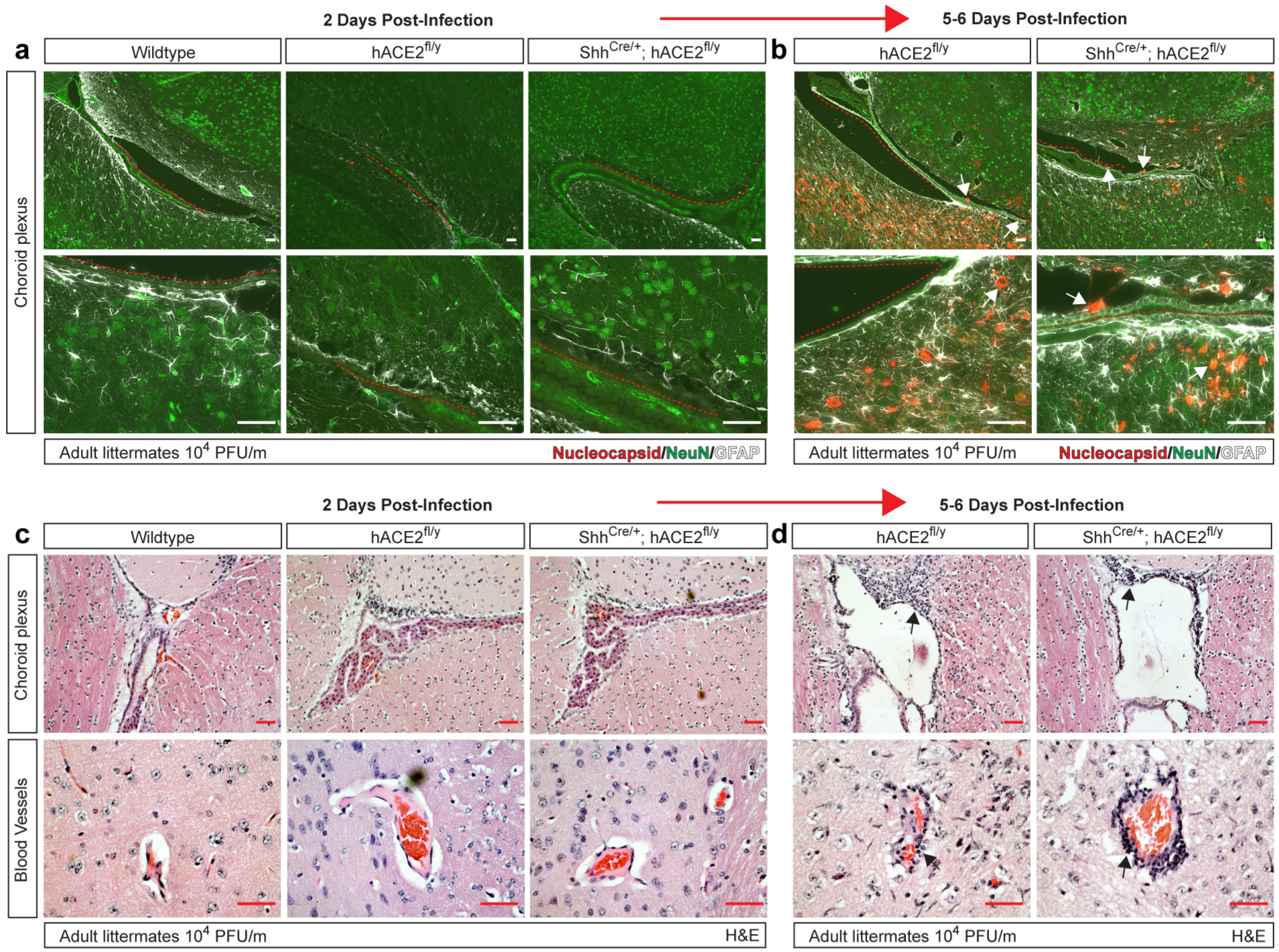
Meningeal and vascular inflammation is associated with neuronal infection at the choroid plexus. **a-b**, Immunohistochemistry of SARS-CoV-2 nucleocapsid, neuronal NeuN and glial cell GFAP of the cerebral cortex adjacent to the choroid plexus 2 and 5-6 days after infection. Arrows indicated nucleocapsid staining. Red dotted lines trace ependymal cells of the choroid plexus. **c-d,** H&E staining of brain choroid plexus and cerebral cortex blood vessels 2 and 5-6 days after infection of *hACE2*^fl/y^ and Shh^Cre/+^; *hACE2*^fl/y^ mice. Arrows indicate sites of immune cell infiltration. Representative of N=4 animals per genotype and timepoint. Scale bars 50 µm.

### Reactive meningitis and encephalitis with adjacent SARS-CoV-2 infection at the choroid plexus

The ciliated epithelium of the choroid plexus expresses genes required for SARS-CoV-2 entry, and a recent study reported an anti-viral inflammatory response at that site in post-mortem biopsies from 8 COVID-19 patients even though no virus could be detected in those samples^33, 34^ (discussed further below). Since our studies of the OE and OB suggested that neuronal infection may be linked to adjacent epithelial cell infection, we next assessed the choroid plexus of infected hACE2 mice. Unlike the OE, SARS-CoV-2 nucleocapsid was not detected in the choroid plexus or adjacent neurons at an early timepoint, 2 days, after infection (Fig. 6a). However, patches of neuronal SARS-CoV-2 nucleocapsid staining were observed 5 days after viral infection in proximity to the choroid plexus and meninges in both *hACE2*^fl/y^ and Shh^Cre^; *hACE2*^fl/y^ animals (Fig. 6b). H&E staining of the choroid plexus along the lateral ventricle and cerebral cortex vasculature revealed the presence of immune cell infiltrates at 5 (but not 2) days after SARS-CoV-2 infection (Fig. 6c, d) indicative of an inflammatory response like that described in humans^34, 35^. These studies suggest that brain inflammation around the choroid plexus may arise due to viral infection at that site at a later timepoint during COVID-19 pathogenesis. Since the patches of infected choroid plexus and adjacent brain were not directly continguous with sites of olfactory bulb and cerebral cortex infection, these studies raise the possibility that more than one route of central nervous system infection may exist.

### Olfactory sensory neurons are infected alongside olfactory epithelium early in SARS-CoV-2 infection

The studies described above, and prior studies of SARS-CoV-2 in the tissues of patients who died of COVID-19 disease^36^, support a pathogenic mechanism in which SARS-CoV-2 virus first infects the OE and subsequently spreads to the brain. Since OSNs are the primary cellular connection between the brain and the ACE2-rich OE that is infected soon after exposure to SARS-CoV-2 (Fig. 5a), we next assessed whether and when OSN infection might take place following viral exposure. 2 days after infection with SARS-CoV-2 virus, *hACE2*^fl/y^ mice exhibited viral RNA in the RE and OE (Fig. 7a). Co-staining for the neuronal marker neurofilament-200 (NF-200, aka NFH) identified penetrating OSNs in the OE that were positive for SARS-CoV-2 RNA (Fig. 7b). 6 days after infection, a timepoint at which infection has progressed to the OB and cerebral cortex, we detected β3-tubulin-labeled, immature OSNs in the OE that co-stained with SARS-CoV-2 nucleocapsid (Fig. 7c). These findings suggest a mechanism in which infection of the OE and OSNs provides viral passage to neurons in the brain.

**Figure 7.**
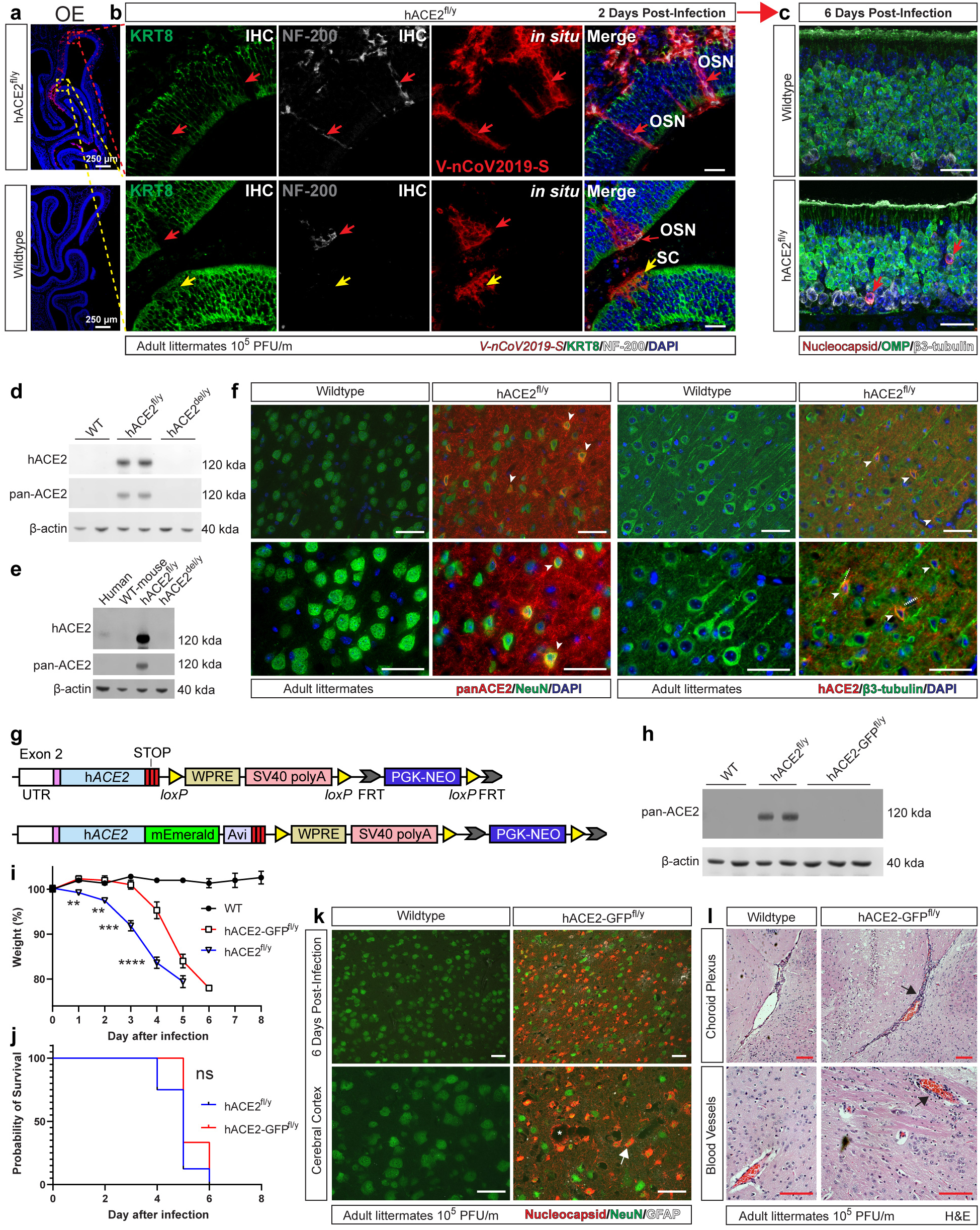
Early infection of olfactory sensory neurons by SARS-CoV-2 and lethal infection of *hACE2-GFP^fl/y^* mice that express low levels of neuronal hACE2. **a**, *In situ* hybridization of SARS-CoV-2 mRNA 2 days post-infection in the olfactory epithelium (OE) of *hACE2*^fl/y^ and wild-type mice infected with 10^5^ titer. b, Higher magnification images of *in situ* hybridization of SARS-CoV-2 mRNA 2 days post-infection in KRT8+ OE sustentacular cells and NF-200+ olfactory sensory neurons of *hACE2*^fl/y^ mice. KRT8, keratin 8; NF-200, neurofilament 200; OSN, olfactory sensory neuron; SC, sustentacular cell. Arrows indicate site of OSNs with colocalized viral mRNA in the olfactory epithelium. Representative of N=3 per genotype. Scale bars 25 µm. c, Immunohistochemistry of wildtype and *hACE2*^fl/y^ olfactory epithelium, 6-days post infection with 10^5^ titer, for SARS-CoV-2 nucleocapsid, olfactory marker protein (OMP, mature OSNs), and βIII-tubulin (immature OSNs). Arrows indicate immature OSNs with colocalized nucleocapsid staining. Representative of N=4 per genotype. Scale bars 25 µm d-e, Immunoblotting of whole brain lysates from human, wild-type mouse, *hACE2*^fl/y^, and *hACE2*^del/y^ animals with antibodies that recognize human and mouse ACE2 (pan-ACE2), or only human ACE2 (hACE2). β-actin antibody concurrently used as a loading control. Representative of N=4 per genotype. f, Immunohistochemistry of ACE2 using pan-ACE2 or human-ACE2 specific antibodies and neurons using NeuN or βIII-tubulin antibodies in wild-type and *hACE2*^fl/y^ mouse cerebral cortex. Arrows indicate NeuN or βIII-tubulin positive neurons colocalized with ACE2 staining. Representative of N=3 per genotype. Scale bars 50 µm. g, Generation of the *hACE2-GFP*^fl/y^ allele using gene targeting of the mouse *Ace2* locus. Monomeric Emerald (mEmerald) Woodchuck hepatitis virus post-transcriptional regulatory element (WPRE), Simian virus 40 (SV40), Phosphoglycerate kinase promoter driven Neomycin resistance cassette (PGK-NeoR). Note: targeted allele is identical to *hACE2*^fl/y^ except for fusion of mEmerald to the c-terminus of hACE2. h, Immunoblotting of whole brain lysates from wild-type (WT), *hACE2*^fl/y^, and *hACE2-GFP*^fl/y^ animals using ACE2 antibodies that recognize both mouse and human ACE2 (pan-ACE2). Representative of N=4 per genotype. i-j, Weight loss and survival of wild-type (WT), *hACE2*^fl/y^, and *hACE2-GFP*^fl/y^ mice after infection with 10^5^ PFU of SARS-CoV-2. Asterisks indicate significant differences in weight between infected *hACE2*^fl/y^, and *hACE2-GFP*^fl/y^ animals. N=6 (WT), 3 (*hACE2-GFP*^fl/y^) and 8 (*hACE2*^fl/y^). **p<0.01; ***p<0.001. ****p<0.0001 by unpaired, two-tailed t-test, ns p>0.05 by log rank Mantel Cox test. Note: the data for *hACE2*^fl/y^ animals same as Figure 2d-e. k, Immunohistochemistry of SARS-CoV-2 nucleocapsid, neuronal NeuN and glial cell GFAP in the cerebral cortex 5 days after infection of wild-type (WT) and *hACE2-GFP*^fl/y^ mice. Representative of N=3 per genotype. Arrows indicate SARS-CoV-2 nucleocapsid-positive neurons. Scale bars, 50 µm. l, H&E staining of wildtype and *hACE2-GFP*^fl/y^ animals infected with 10^5^ viral titer harvested 6 days post-infection. Shown are meninges along the lateral ventricle (top row) and cerebral cortex blood vessels (bottom row). Arrows indicate perivascular immune cell infiltrate. N=3 per genotype, one experiment. Scale bars 50 µm.

To further investigate the mechanism of SARS-CoV-2 brain infection in *hACE2*^fl/y^ mice, we compared the expression of ACE2 in wild-type and *hACE2*^fl/y^ mouse brains. Immunostaining of brain sections and immunoblotting of brain cell lysate using pan-ACE2 antibodies that recognize both mouse and human ACE2 protein revealed strong expression of ACE2 in *hACE2*^fl/y^ brains (Fig 7d-f). In contrast, ACE2 could not be detected using either method in wild-type mouse brain (Fig. 7d-f) and was detected at lower levels in human brain cell lysate (Fig. 7e). These results suggested that *hACE2*^fl/y^ mice express levels of ACE2 in the brain higher than those in wild-type mice and humans, raising the possibility that brain infection may be a consequence of elevated levels of brain ACE2 in that model and not accurately reflect infection in the wild-type mouse or human brain.

To address this possibility, we generated a second line of ATG-knock in animals: *hACE2-GFP*^fl^ mice, in which the hACE2 protein is fused to monomeric Emerald (mEmerald, a green fluorescent protein (GFP) mutated to prevent auto-dimerization and improve solubility) using the same conditional strategy previously described for *hACE2*^fl^ mice (Fig. 7g). Immunoblotting of kidney cell lysate and immunostaining of small intestine tissue sections demonstrated appropriate expression of the hACE2-GFP protein in ciliated epithelial cells in vivo (Ext. Data Fig. 7). In contrast to *hACE2*^fl/y^ brains, but like those of wild-type mice and healthy humans, *hACE2-*GFP^fl/y^ brains expressed undetectable levels of hACE2 protein using immunoblotting of brain cell lysate (Fig. 7h). To test whether brain infection by SARS-CoV-2 was merely a consequence of high brain ACE2 expression in *hACE2*^fl/y^ mice, we next infected *hACE2-GFP*^fl/y^ mice with SARS-CoV-2 virus. *hACE2-GFP*^fl/y^ mice exhibited weight loss 4 days after infection and lethality by day 6 (Fig. 7i, j). Significantly, analysis of the cerebral cortex 6 days after infection revealed SARS-CoV-2 nucleocapsid in NeuN-positive neurons and reactive gliosis of *hACE2-GFP*^fl/y^ animals (Fig. 7k), consistent with neuronal infection like that observed in *hACE2*^fl/y^ mice. H&E staining also revealed meningeal and peri-vascular inflammatory infiltrates (Fig. 7l), as well as patchy peri-meningeal sites of neuronal nucleocapsid staining like those seen in *hACE2^fl/y^* mice (Ext. Data Fig. 8). These studies confirm that neuronal infection takes place in animals that express levels of hACE2 in the brain like those in the wild-type mouse and human brain and provide additional evidence for a pathogenic link between infection of the OE, OSN and brain during lethal COVID-19.

**Figure 8.**
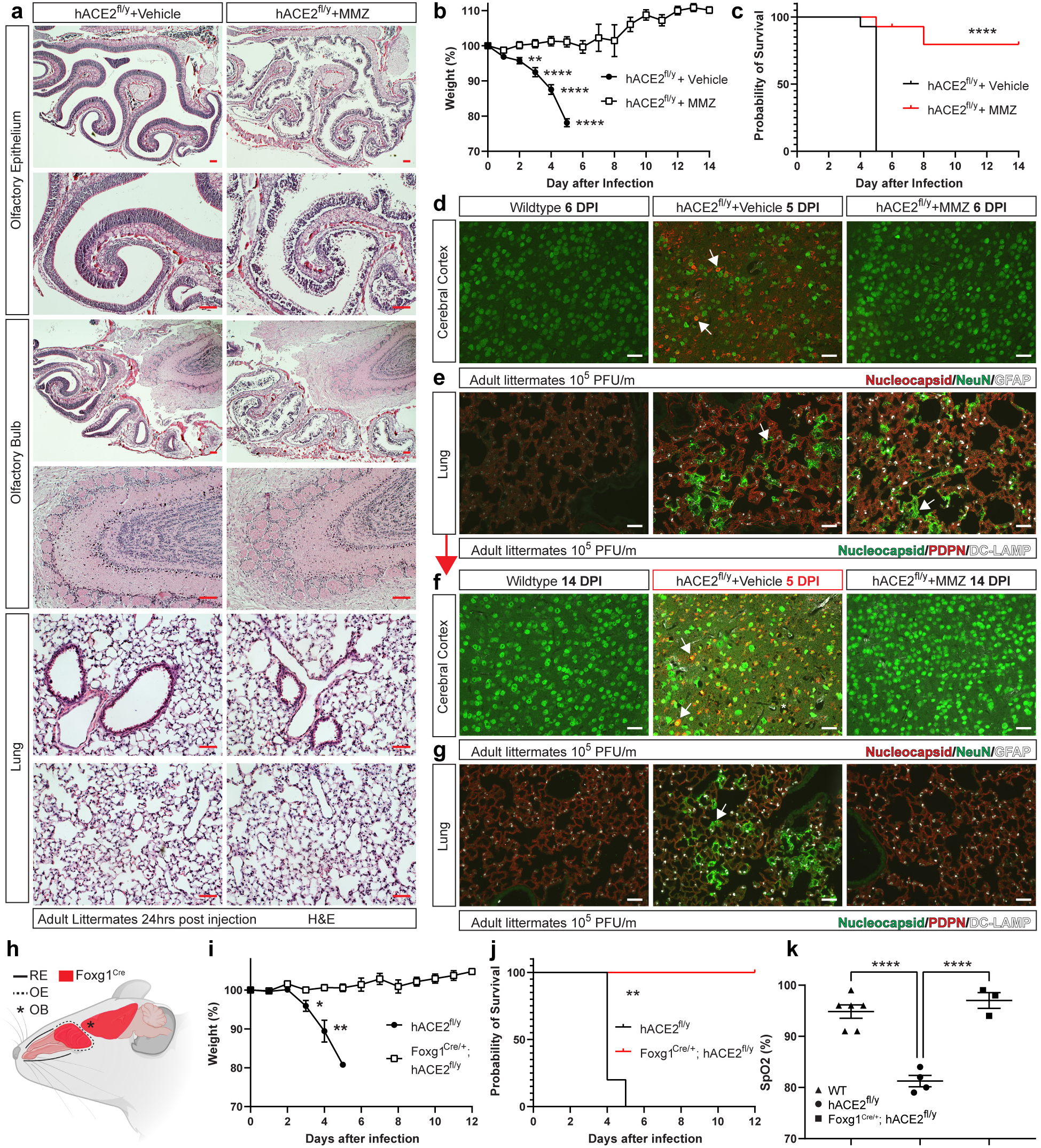
Methimazole ablation of the OE or genetic loss of hACE2 in OE and neurons prevents brain infection and lethal SARS-CoV-2 infection. **a**, H&E staining of the indicated mouse tissues was performed 24 hours after intraperitoneal injection of 100 mg/kg methimazole (MMZ) or vehicle control. Scale bars 100 µm for the olfactory epithelium, olfactory bulb and 50 µm for the lung. Representative of N=3 per condition. b-c, Weight loss and survival of *hACE2*^fl/y^ mice treated with MMZ or vehicle 24 hours prior to infection with 10^5^ PFU of SARS-CoV-2 virus. Asterisks indicate significant differences in weight between vehicle and MMZ treated *hACE2^fl/y^* animals. N=14 (Vehicle) and 15 (MMZ). d-e, Immunohistochemistry of wild-type, vehicle-treated *hACE2*^fl/y^ and MMZ-treated *hACE2*^fl/y^ mouse cerebral cortex and lung 5-6 days after infection with 10^5^ PFU of SARS-CoV-2 virus using antibodies that recognize viral nucleocapsid, the neuronal marker NeuN, the glial cell marker GFAP, the alveolar type II cell marker DC-LAMP, the alveolar type I (AT1) cell marker PDPN and the nuclear stain DAPI. Arrows indicate nucleocapsid staining colocalized with NeuN+ neurons (brain) or PDPN+ AT1 cells (lung). Representative of N=4 per condition. f-g, Immunohistochemistry of wild-type and MMZ-treated *hACE2*^fl/y^ mouse cerebral cortex and lung 14 days after infection with 10^5^ PFU of SARS-CoV-2 virus was performed as described in d-e. Note: Day 5 *hACE2*^fl/y^ samples (middle panel) were included on the same tissue slide as a positive control. Arrows indicate nucleocapsid staining colocalized with neurons and AT1 cells in vehicle-treated *hACE2*^fl/y^ mice. Representative of N=4 per condition. Scale bars d-g 50 µm. h, The Foxg1^Cre/+^ allele drives Cre expression in the olfactory epithelium (OE) and neurons of the brain (shown in red). i-j, Weight loss and survival of *hACE2*^fl/y^ and and Foxg1^Cre/+^; *hACE2*^fl/y^ mice after infection with 10^5^ viral titer per mouse. n=4 and 3 mice, respectively. k, Pulse oximetry measured in wild-type (WT), *hACE2*^fl/y^ and Foxg1^Cre/+^; *hACE2*^fl/y^ mice 5-6 days after exposure to SARS-CoV-2 virus. *p<0.05, ** p<0.001; **** p<0.0001 by unpaired two-tailed t-test, one-way ANOVA with Holm-Sidak correction for multiple comparisons, or log-rank Mantel Cox test.

### Pharmacologic ablation of the OE or genetic loss of hACE2 in the OE and neurons prevents lethal COVID-19 disease

The studies described above indicated that SARS-CoV-2 infection of the OE rather than the lung is associated with weight loss and lethality. Since genetic tools to drive Cre expression selectively in the OE are not available, we tested the requirement for OE infection using pharmacologic ablation of olfactory epithelial cells with methimazole (MMZ), a compound that is highly and specifically toxic for the OE in rodents^37^. H&E staining of the OE 24 hours after treatment of *hACE2*^fl/y^ mice with 100 mg/kg methimazole revealed loss of almost all olfactory epithelial cells (Fig. 8a). In contrast, the underlying olfactory bulb and lung did not show evidence of cell loss, damage, or inflammation (Fig. 8a). We next tested whether ablation of the OE with MMZ impacts the clinical course of *hACE2*^fl/y^ mice infected with SARS-CoV-2 virus. Remarkably, pre-treatment with MMZ 24 hours prior to intranasal inoculation of SARS-CoV-2 prevented both weight loss and death in the majority of *hACE2^fl/Y^* mice (Fig. 8b, c).

Immunostaining of the brain for SARS-CoV-2 nucleocapsid protein at 6 days after infection revealed the absence of detectable virus in the cerebral cortex of MMZ-treated *hACE2*^fl/y^ mice 6 days after infection compared with vehicle-treated controls (Fig. 8d). In contrast, similar expression of SARS-CoV-2 nucleocapsid was detected in the lungs of vehicle-treated and MMZ-treated *hACE2*^fl/y^ mice (Fig. 8e). Analysis of MMZ-treated *hACE2*^fl/y^ mice 14 days after SARS-CoV-2 infection revealed no viral nucleocapsid in either the brain or lung (Fig. 8f, g), consistent with sparing of the brain and resolution of lung infection in surviving MMZ-treated *hACE2*^fl/y^ mice. Consistent with brain infection being a driver of lethality, analysis of a single *hACE2*^fl/y^ mouse that became ill and was euthanized 8 days after infection revealed strong staining for nucleocapsid in the brain and lung (Ext. Data Fig. 9).

**Figure 9.**
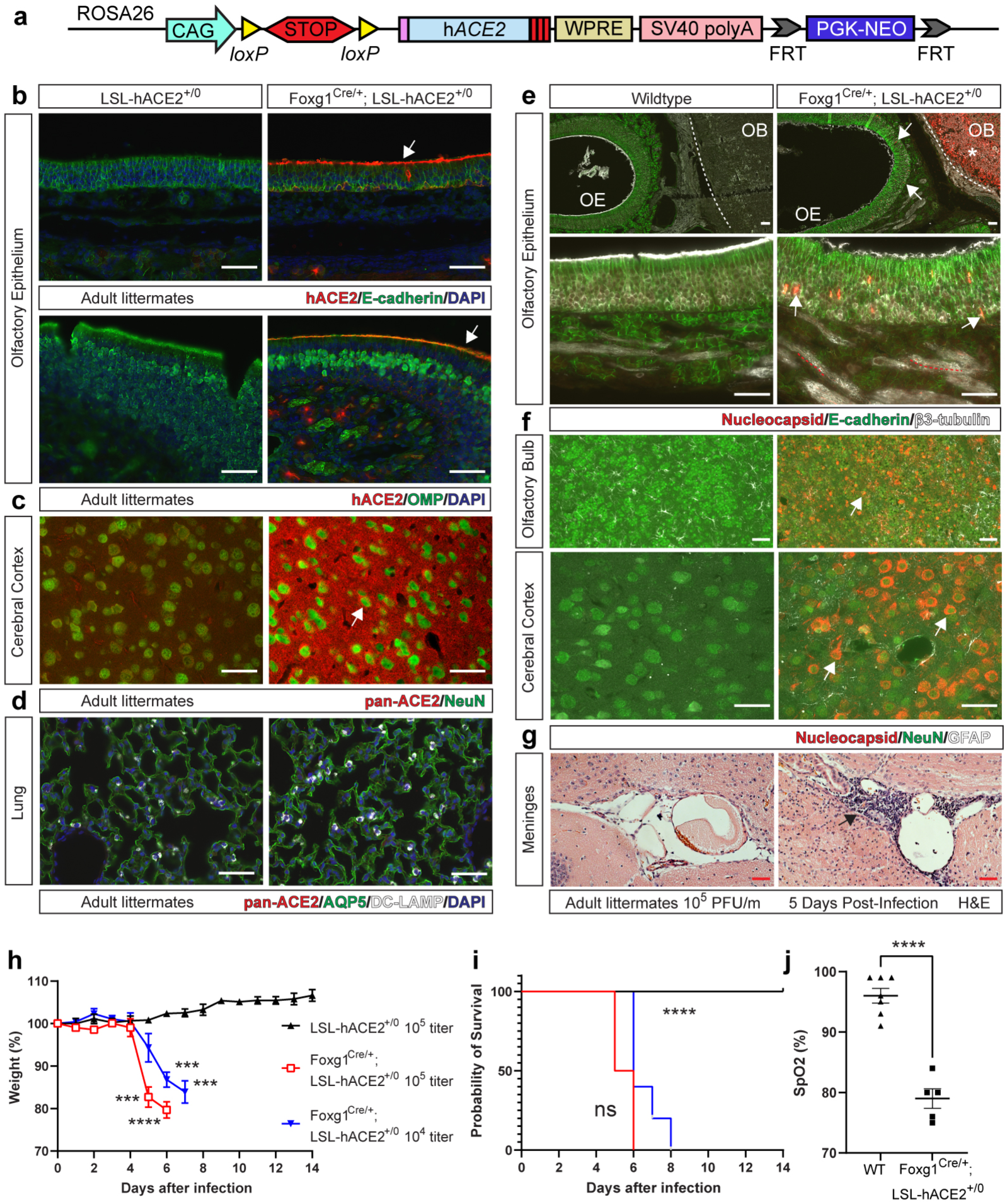
Forced expression of hACE2 in OE and neurons is sufficient to confer hypoxia and death following SARS-CoV-2 infection. **a**, Generation of LSL-hACE2 mice using gene targeting of the mouse *Rosa26* locus. Woodchuck hepatitis virus post-transcriptional regulatory element (WPRE), Simian virus 40 (SV40), Phosphoglycerate kinase promoter driven Neomycin resistance cassette (PGK-NeoR). **b-d**, Immunohistochemistry of olfactory epithelium, cerebral cortex and lung from Foxg1^Cre/+^;LSL-hACE2^+/0^ and LSL-hACE2^+/0^ mouse brain tissue was performed using antibodies that recognize human ACE2, (hACE2), both human and mouse ACE2 (pan-ACE2), the neuronal marker NeuN, alveolar type 1 cell Aquaporin 5 (AQP5), and alveolar type II cells (DC-LAMP). Representative of N=2 per genotype. Scale bars 50 µm. **e-f**, Immunohistochemistry of Foxg1^Cre/+^;LSL-hACE2^+/0^ and LSL-hACE2^+/0^ mouse olfactory epithelium, olfactory bulb, and cerebral cortex 5-6 days after infection with 10^5^ PFU of SARS-CoV-2 virus using antibodies that recognize viral nucleocapsid, sustentacular cell E-cadherin, olfactory sensory neuron (OSN) βIII-tubulin, neuronal NeuN, or glial GFAP. OE-olfactory epithelium. OB-olfactory bulb. Asterisk indicates olfactory bulb with nucleocapsid staining. Arrows indicate nucleocapsid staining colocalized with βIII-tubulin+ OSNs (olfactory epithelium) or NeuN+ neurons (brain). Red dotted lines trace nucleocapsid staining colocalized within β3-tubulin+ OSN axons. Representative of N=4 per genotype. Scale bars 50 µm. **g-h**, Weight loss and survival of LSL-hACE2^+/0^ and Foxg1^Cre/+^; LSL-hACE2^+/0^ mice after infection with 10^5^ or 10^4^ PFU of SARS-CoV-2. Asterisks indicate timepoints at which significant differences in weight were observed between infected Foxg1^Cre/+^; LSL-hACE2^+/0^ and LSL-hACE2^+/0^ mice animals. *** P<0.001; **** P<0.0001 determined by unpaired, two-tailed t-test or log-rank Mantel Cox test. **i**, Pulse oximetry of WT (LSL-hACE2^+/0^ or Foxg1^Cre/+^) and Foxg1^Cre/+^; LSL-hACE2^+/0^ mice infected with 10^4^ viral titer at the time of harvest (Day 6-8). ****p<0.0001 determined by unpaired, two-tailed t-test.

To further test the requirement for SARS-CoV-2 infection of the OE and brain for lethal COVID-19 in *hACE2*^fl/y^ mice, we generated Foxg1^Cre/+^; *hACE2*^fl/y^ mice in which Cre recombinase is expressed in the OE and neurons of the forebrain, but not in cells of the lung^38^ (Fig. 8h). Following exposure to SARS-CoV-2, *hACE2^fl/y^* littermate mice exhibited weight loss and lethality associated with severe hypoxia, like previous experiments in Figures 2 and 4. In contrast, Foxg1^Cre/+^; *hACE2*^fl/y^ mice survived without weight loss or hypoxia (Fig. 8j, k). Following exposure to SARS-CoV-2, *hACE2^fl/Y^* littermate mice exhibited weight loss and lethality associated with severe hypoxia, as in previous experiments shown in Figures 2 and 4. In contrast, *Foxg1^Cre/+^*; *hACE2^fl/y^* mice survived and did not display weight loss or hypoxia (Fig. 8j, k). These pharmacologic and genetic findings demonstrate that SARS-CoV-2 infection of the OE is required for brain infection and lethal COVID-19 in *hACE2*^fl/y^ mice, while infection of the lung is not sufficient to cause weight loss and death.

### SARS-CoV-2 infection of the olfactory epithelium and brain is sufficient to confer acute lung injury and lethal COVID-19

The above studies demonstrated that ACE2-dependent viral infection is required in the OE but not in the lung for decompensation following exposure to SARS-CoV-2, and identified a strong association between the OE, neuronal infection, and lethality in all hACE2 mouse models studied. Thus, infection of olfactory epithelial cells and neurons is either sufficient to confer severe COVID-19 or is required for infection of another, yet unidentified, cell population. To distinguish between these possibilities and rigorously test the sufficiency of OE and neuronal infection for severe disease, we generated a gain-of-function model in which Cre recombinase drives cell-specific expression of hACE2 from the *Rosa26* locus (“LSL-hACE2 mice”, Fig. 9a). LSL-hACE2 mice were crossed to Foxg1^Cre/+^ animals to generate Foxg1^Cre/+^; LSL-hACE2^+/0^ mice in which hACE2 is expressed exclusively in the OE and neuronal cells. Immunostaining using anti-hACE2 or anti-pan-ACE2 antibodies revealed expression of hACE2 in the OE and brain of Foxg1^Cre/+^; LSL-hACE2^+/0^ animals but not LSL-hACE2^+/0^ littermates (Fig. 9b, c), consistent with Cre-dependent hACE2 expression. In contrast, hACE2 could not be detected in the lung of either Foxg1^Cre/+^;LSL-hACE2^+/0^ or control LSL-hACE2^+/0^ littermates (Fig. 9d), consistent with previously reported lineage tracing^38^. Exposure of Foxg1^Cre/+^;LSL-hACE2^+/0^ and LSL-hACE2^+/0^ mice to SARS-CoV-2 resulted in viral infection of the nasal OE and brain but not the lung (Fig. 9e,f and Ext. Data Fig. 10). Similar to findings in *hACE2*^fl/y^ and *hACE2-GFP*^fl/y^ mice, an immune cell infiltrate was observed adjacent to the choroid plexus and meninges of the lateral ventricle 5 days after infection (Fig. 9g). SARS-CoV-2 infection caused weight loss, hypoxemia and death in Foxg1^Cre/+^;LSL-hACE2^+/0^ animals, with no evidence of illness in LSL-hACE2^+/0^ littermates (Fig. 9h-j).

Severe hypoxemia associated with lethal SARS-CoV-2 infection in Foxg1^Cre/+^;LSL-hACE2^+/0^ animals suggested that viral infection of the OE and brain might be sufficient to confer pulmonary disease like that observed in *hACE2*^fl/y^ mice. Indeed, histologic analysis of the lungs of infected Foxg1^Cre/+^;LSL-hACE2 animals revealed lung epithelial inflammatory changes and intravascular thrombosis similar to those seen in infected *hACE2*^fl/y^ mice (Fig. 4 and Ext. Data Fig. 4) despite the lack of lung infection (Ext. Data Fig. 10). These findings demonstrate that SARS-CoV-2 infection of the olfactory epithelial and neuronal cell populations is sufficient to confer severe COVID-19 associated with an acute lung injury/ARDS phenotype in mice— findings highly consistent with the loss of function studies described above (Fig. 8).

## Discussion

Rigorous testing of COVID-19 disease mechanisms has been impeded by the lack of an animal model that permits genetic dissection of tissue- and cell type-specific requirements for pathogenesis. Our studies utilize new mouse genetic models that enable Cre-mediated loss and gain of human ACE2 expression to confer severe COVID-19 and examine this important question. We find that lung infection by SARS-CoV-2 is not required for lethality, acute lung injury, and hypoxemia in mice. Instead, genetic loss of function, genetic gain of function, and pharmacologic studies support a central role for the olfactory epithelium (OE) and neighboring neuronal cells in the pathogenesis of severe COVID-19. These studies shed new light on the cellular mechanisms underlying severe SARS-CoV-2 infection and suggest that strategies targeting the OE and brain may be valuable approaches towards therapeutics and infection prophylaxis.

Diffuse organ involvement has been documented following SARS-CoV-2 infection, but the most dangerous presentation is one in which patients develop rapidly progressive hypoxemia associated with alveolar filling and acute respiratory distress syndrome (ARDS)^13, 30^. While respiratory failure associated with ARDS can arise due to extra-pulmonary etiologies, most are associated with pulmonary infection and lung injury^39^. Thus, it has been assumed that primary viral infection of the lung, most likely involving the AT2 cell in which low ACE2 expression is detectable, is a causal event. Our studies identify SARS-CoV-2 virus in the lungs of infected hACE2 mice but demonstrate that AT1 cells are also infected despite the inability by us and others to detect ACE2 in AT1 cells in the lung^3, 26^. These results are also consistent with temporally controlled experiments in primates^40, 41^. Importantly, AT1 cell infection is ACE2-dependent as we observed complete loss of SARS-CoV-2 in the lungs of Shh^Cre^; *hACE2*^fl/y^ animals. However, despite a lack of lung infection, Shh^Cre^; *hACE2*^fl/y^ animals exhibited hypoxemia and weight loss comparable to those observed in *hACE2*^fl/y^ littermates after infection with SARS-CoV-2. Taken together, these studies highlight the value of functional genetic approaches to dissecting disease mechanism at the cellular level and reveal (i) that the level of cellular ACE2 expression required for SARS-CoV-2 infection is below that of detection using standard methods such as immunostaining, in situ hybridization and single cell RNAseq, and (ii) that a lethal COVID-19 ARDS phenotype may arise due to extra-pulmonary infection. The extent to which pulmonary vs extra-pulmonary mechanisms of respiratory failure participate in disease is likely to vary between patients and be influenced by individual co-morbidities and immune responses. In addition, the importance of lung epithelial infection for tissue repair and regeneration in the convalescent setting remains unknown.

The most surprising finding in this study is that SARS-CoV-2 infection of OE is both required and sufficient for severe disease in mice. Our studies support a pathogenic mechanism in which inhaled SARS-CoV-2 virus infects sustentacular cells expressing ACE2 protein localized to microvilli that project into the nasal cavity. Olfactory sensory neurons (OSNs) adjacent to sustentacular cells are infected at the same early timepoint (2 days after infection), while neurons in adjacent regions of the brain demonstrate viral infection at later timepoints. Ablation of the OE with methimazole (MMZ) treatment or genetic deletion of hACE2 in the OE protects against weight loss and death, while gain of hACE2 expression specifically in olfactory epithelial cells and cerebral cortex neurons is sufficient to confer lethality and acute lung injury. These results are consistent with prior studies identifying brain infection in human patients^36, 42, 43^, mouse ^7, 15, 21, 44, 45^ and hamster^32^ models, and *ex vivo* organoid systems^6, 44, 46, 47^ and support a mechanism in which infection of the OE enables infection of central nervous system neurons. This pathogenesis provides a straightforward explanation for the anosmia and persistent neurologic symptoms such as “brain fog” that are characteristic of human COVID-19^48–50^.

However, SARS-CoV-2 neuronal infection has been refuted by prominent studies that failed to identify virus in the brains of deceased patients using methods such as single nuclear RNA sequencing^34^ or RNA *in situ* hybridization and immunostaining of olfactory bulb neurons^51^. If infection of the OE and neighboring brain are associated with lethal COVID-19 disease, why is SARS-CoV-2 virus detected in human brain samples by some but not all investigators? We find that the virus moves rapidly from one tissue to another in a patchy fashion, suggesting that there may be a narrow temporal and spatial window in which to detect SARS-CoV-2 in a particular cell type. Examination of post-mortem brains from patients whose disease course has been extended by aggressive treatment such as mechanical ventilation (e.g. greater than 2 weeks^34^) may bias towards analysis after virus is cleared. Human biopsies are also limited in their sample size and may miss important sites of infection—an explanation for negative findings supported by the fact that SARS-CoV-2 could not be detected in nasal epithelium or brain in more than half of individuals examined by a recent negative study^51^. In contrast, thorough examination of whole nasal cavity and brain from mice infected with a measured amount of virus at a known timepoints during a highly reproducible disease course enables detection of even transiently infected tissues.

Our findings have implications for treatments and vaccines directed against SARS-CoV-2. Of immediate relevance is assessing the bioavailability and pharmacodynamics of current and future therapies in the central nervous system. While design of drugs that overcome the blood brain barrier is challenging, the ability of MMZ to protect against lethal, acute disease in hACE2 mice suggests that preventing infection in the OE may be an efficient and targeted means of protecting against severe disease. Importantly, MMZ itself is likely not a viable treatment for COVID-19 as it is an anti-thyroid medication and the high doses required to ablate olfactory epithelium in rodents have not been studied in humans. Instead, vaccines aimed at stimulating mucosal immunity or agents that interfere with SARS-CoV-2 binding to ACE2 at the OE administered via a nasal spray may confer long or short-term protection without systemic side effects. Such nasal prophylactics could be rapidly altered to protect against new viral variants and be considered less dangerous or more acceptable to vaccine-hesitant individuals.

Interpretation of this study must consider several limitations. First, the mouse OE is more developed compared to human OE; thus, a mouse model may bias toward disease mechanisms involving the OE. In this regard, it is notable that we see complete lethality after exposure to virus in hACE2 mice while most humans experience mild disease. This outcome may also reflecs the administration of high viral titers that greatly exceed the inoculum received by most human patients. Second, since mice are obligate nose-breathers, our studies utilize intranasal SARS-CoV-2 administration. Unlike mice, humans are capable of mouth-breathing; thus, transmission may be less dependent on the OE. However, anosmia is pathognomonic for human disease and often precedes the onset of respiratory failure—a sequence consistent with that we observed in hACE2 mice. Overall, our mouse models likely reflect the most severe disease course experienced by human patients, e.g. those with the largest viral inocula and/or the greatest number of preexisting conditions that act as disease risk factors.

Studies of human patients that more directly examine infection of the OE and brain early in disease are challenging but will be required to fully validate and extend these findings. In addition, the molecular mechanism by which SARS-CoV-2 infection of olfactory epithelial cells and neurons leads to acute lung injury in hACE2 mice is an important future direction with clear therapeutic implications. Finally, our studies have exclusively examined acute responses to SARS-CoV-2 infection. Optimization of this hACE2 model to enable interrogation of long-term sequelae after recovery from acute infection are likely to elucidate mechanisms that underlie other important aspects of human COVID-19.

## Acknowledgements

We thank the members of the Kahn lab for their thoughtful comments and advice during this work. We thank the CDB Microscopy Core for microscopy support. Schematics of the mouse brain and respiratory tract were created with BioRender.com.

## Funding

Supported by National Institute of Health grants R01H139552-04S1 (MK), R01AI109022 and R21AI156731 (HAC), T32EB023860 (DWB), R25GM125597 (BI), and R01AI138370 (BI and AA), and a Penn Cardiovascular Institute Dream Team grant (MK).

## Author Contributions

ATT designed and targeted the hACE2 mouse lines used in this manuscript. ATT and DWB designed and performed most of the mouse studies and wrote the manuscript. KMS, SG, MF, MW, JY, PM-I, XZ, and MX performed immunostaining and immunoblotting studies. PH and IAM generated the virus used for infections. BI and PH performed mouse SARS-CoV-2 infection studies. JS and PM-I assisted with mouse handling and transfer. NAL and SS performed ES cell microinjections to generate chimeric F0 mice. AA provided funding and mentoring. KJ, MX, EEM, SEM, HCA and MLK designed experiments and wrote the manuscript.

## Competing interests

The authors declare no competing financial interests.

## Data and materials availability

All data and reagents will be made available upon reasonable request. Transgenic mouse lines not available through public repositories are available from Mark Kahn under a material transfer agreement with the University of Pennsylvania.

## Materials and Methods

### Mice

The humanized ACE2 (hACE2) floxed and hACE2-GFP floxed alleles were generated using CRISPR/Cas9 assisted mouse embryonic stem cell targeting as previously described^52^. A gRNA sequence 5’-GGATGGGATCTTGGCGCACG-3’ targeting intronic sequence immediately prior to Exon 2 of *Ace2* was cloned into the eSpCas9(1.1) plasmid (Addgene 71814). A targeting construct was designed to insert by homologous recombination the human *ACE2* cDNA followed by a floxed WPRE-SV40 polyA and FRT-ed neomycin resistance cassette directly after the ATG-start codon of mouse *Ace2* in exon 2. The human ACE2 cDNA sequence was modified to utilize the mouse ACE2 signal peptide, codon optimized for mouse codon usage, then further altered with silent mutations to remove cryptic RNA splice sites. The same hACE2 targeting construct was utilized to generate the hACE2-GFP floxed targeting construct except with the addition of a monomeric-Emerald GFP protein fused to the c-terminal sequence of hACE2 through a GSG-flexible linker sequence. Both targeting constructs were synthesized by Genscript. Properly targeted ES cell clones were confirmed using PCR screening of the 5’ and 3’ arms of homology and the entire targeted knock-in was PCR amplified for Sanger sequencing prior to microinjection into V6.5 ES cells. Chimeras were mated to B6D2F1/J (Jackson Laboratory, 100006) females to generate germline F1 mice. Genotyping primers that distinguish zygosity against the 5’ region of the knock-in are as follows: wildtype Forward 5’-ctcagtgcccaacccaagttc-3’, wildtype Reverse 5’-atgtcttggcattttcctcggt-3’, and mutant Reverse 5’-ggagctggagctttacggtga-3’ with expected band sizes of 400 bp (mutant) and 190 bp (wildtype).

The Rosa26-LSL-hACE2 allele was also generated using CRISPR/Cas9 assisted mouse embryonic stem cell targeting. A gRNA sequence 5’-cgtgatctgcaactccagtc-3’ targeting the XbaI restriction site conventionally used for Rosa26 targeted alleles was cloned into the eSpCas9(1.1) plasmid (Addgene 71814). The targeting construct was synthesized by Genscript and screening of targeted ES cell clones was performed as aforementioned. Genotyping primers against the 5’ region of the knock in that distinguish zygosity are as follows: wildtype Forward 5’-ttctgggagttctctgctgc-3’, wildtype Reverse 5’-tgggaagtcttgtccctcca-3’, and mutant Reverse 5’-agagtgaagcagaacgtggg-3’ with expected band sizes of 423 bp (mutant) and 210 bp (wildtype).

Shh-Cre (stock 005622), Sox2-Cre (stock 008454), Sftpc-CreERT2 (stock 028054), R26-LSL-tdTomato (Ai14, stock 007914) animals were purchased from Jackson Laboratories on the congenic C57BL/6J strain^28, 53, 54^. Shh-Cre was inherited from the male in all experiments given the known mosaic expression of Shh-Cre from maternal inheritance. Sox2-Cre was used as a germline deleter via maternal inheritance to establish a germline recombined allele. The Foxg1-IRES-Cre was a kind gift from Nada Jabado^38^.

Experimental mice were maintained on a mixed C57BL/6J, 129S1/SvJ, and DBA/2J background and utilized between 2-4 months of age for infection studies. Mice were housed in a specific pathogen free facility where cages were changed on a weekly basis. Cages, bedding, food, and acidified water (pH 2.5-3.0) were autoclaved prior to use. Ambient temperature maintained at 23 degrees Celsius, and 5% Clidox-S^TM^ was utilized as disinfectant. The University of Pennsylvania Institutional Animal Care and Use Committee (IACUC) approved all animal protocols, and all procedures were performed in accordance with these protocols (#806811).

### Penn ABSL3 facility and viral inoculation

Animal studies were carried out in accordance with the recommendations in the Guide for the Care and Use of Laboratory Animals of the National Institutes of Health. The protocols were approved by the Institutional Animal Care and Use Committee (IACUC) at the University of Pennsylvania (protocol #807017). Virus inoculations were performed under anesthesia that was induced and maintained with ketamine hydrochloride and xylazine, and all efforts were made to minimize animal suffering. Animals were housed in groups and fed standard chow diets. Mice of different ages and both sexes were administered 1 × 10^5^ plaque forming units (PFU) of SARS-CoV-2 via intranasal administration.

### Penn SARS-CoV-2 Virus

SARS-CoV-2 (Isolate USA-WA1/2020) was obtained from BEI Resources. It was deposited by the Centers for Disease Control and Prevention and obtained through BEI Resources, NIAID, NIH: SARS-Related Coronavirus 2, Isolate USA-WA1/2020, NR-52281. Infectious stocks were grown in Vero-E6 cells and stored at −80 °C. All work with infectious virus was performed in a biosafety level 3 laboratory and approved by the Institutional Biosafety Committee and Environmental Health and Safety.

### Cornell ABSL3 facility and viral inoculations

Mice were anesthetized with isoflurane then intranasally infected with SARS-CoV-2 USA-WA1/2020 (BEI resources; NR-52281) by applying drops of pre-characterized viral stock into the rostral meatus of the nose for a total volume of 50 µL per mouse. Mice were monitored and weighed daily, then euthanized when they lost 20% of their starting weights as a predefined humane endpoint. All mice studies were performed in a BSL-3 laboratory with accordance to protocols approved by the Institutional Animal Care and Use Committee at Cornell University (IACUC mouse protocol # 2017-0108 and BSL3 IBC # MUA-16371-1).

### Viral titers and plaque forming unit (PFU) assays

Vero E6 cells were seeded in 12 well plates with complete DMEM and incubated at 37°C for 24 hours prior to infection. Lung tissue was homogenized in DMEM with 2% serum at 20ul/mg lung tissue. Supernatants were serially diluted and 100ul used to infect cells in duplicates for 1hr at 37°C, gently mixed every 10 minutes, and overlayed with 1 ml of overlay medium (DMEM with 4% serum and 0.3% oxoid agar). The cells were incubated at 37°C for 3 days, fixed with 4% PFA for 30 minutes, then stained with 200ul of 0.5% crystal violet in 30% methanol for 15 minutes. Crystal violet was removed, the cells were washed three times with 1 ml water and plaques counted manually.

### Pulse oximetry

MouseStat Jr X (Kent Scientific) was used to measure heartrate and oxygen saturation (SpO2) of mice prior to harvest. To acclimate the mice to recordings, the sensor was attached daily during routine observation and weighing. Briefly, the sensor was applied to the rear leg and data was recorded following readout stabilization defined as an unchanging recording over 5 seconds.

### Tamoxifen administration

To induce CreERT2 mediated recombination, animals at 8 weeks of age were gavaged with 100 µL of a 30 mg/mL solution of tamoxifen/corn oil (Sigma Aldrich T5648). This was done 3 consecutive days then animals were allowed to rest for 4 days before another three consecutive daily doses (6 doses over 10 days). Animals were used for viral infection studies 2 weeks after completion of the tamoxifen dosing.

### Methimazole administration

A 20 mg/mL solution of methimazole/sterile PBS (Sigma Aldrich M8506) was used to administer a 100 mg/kg dose via intraperitoneal injection. Animals were inoculated with SARS-CoV-2 virus 24-28 hours after methimazole injection.

### Mouse harvest and tissue processing

Mice were euthanized with ketamine/xylazine and underwent laparotomy and sternotomy with subsequent left and right ventricular cardiac perfusion with 20 mL total of PBS. Trachea was exposed and cannulated to allow for bilateral lung inflation. Lungs were gently inflated with PBS infusion, which was stopped upon observation of terminal lung inflation to avoid over-inflation artifacts. Right upper and middle lobes were frozen at -80 degrees Celsius for downstream plaque forming unit assays. Right lower lobe was homogenized in Trizol and stored at -80 degrees Celsius. Left lobe was fixed in 4 percent paraformaldehyde/PBS (volume/volume) along with other organs of interest for a minimum of 72 hours to ensure viral inactivation. At this point, tissues were removed from the animal BSL3 facility and underwent ethanol dehydration and embedding in paraffin blocks for histology.

The intact skull with skin/fur and eyes removed was fixed for a minimum of 72 hours and removed from the animal BSL3 facility. The lower jaw was removed, and nasal cavity was bisected in the sagittal orientation to preserve respiratory and olfactory epithelium. The nasal cavity underwent de-calcification for 14 days in 0.5 M EDTA pH 8.0 at 4 degrees Celsius with agitation prior to ethanol dehydration and paraffin embedding. The brain was removed from the skull and dehydrated with ethanol for subsequent paraffin embedding and histology.

### qPCR for viral loads

Lung tissue was homogenized in Trizol using a bead mill then stored at -80 degrees Celsius. Whole blood was mixed with twice the volume of Trizol LS vortexed then stored at -80 degrees. RNA was extracted using chloroform phase separation and the RNeasy Mini Kit (Qiagen 74004). 1 ug of RNA was reverse transcribed with Superscript IV VILO master mix (Thermo Fisher 11756050) and diluted 1:20 prior to use in qPCR reactions. qPCR was performed using the SARS-CoV-2 Research Use Only qPCR Probe Kit (IDT 10006713) with a standard curve using a positive control N-protein plasmid (IDT 10006625).

### Immunohistochemistry

Paraffin slides with 5 µm thick tissue sections were deparaffinized with xylene and ethanol. Antigen retrieval performed with sodium citrate buffer (Sigma-Aldrich C9999) and a steamer. Primary antibodies were incubated overnight at 4 degrees Celsius and Alexa Fluor secondary antibodies were incubated at room temperature for two hours prior to tissue mounting with Prolong Gold (Thermo Fisher P36930). Importantly, all control and experimental samples were embedded on the same slide and underwent staining under identical conditions. Imaging was performed using identical settings.

Primary antibodies: pan-ACE2 (goat, 1:1000, R&D AF933), human-ACE2 (rabbit, 1:200, Abcam ab108209), E-cadherin (rabbit, 1:200, Cell Signaling 3195S), E-cadherin (goat, 1:200, R&D AF748), Olfactory marker protein (goat,1:200, Wako 544-10001), SARS-CoV-2 nucleocapsid (rabbit, 1:500, Rockland 200-401-A50), Podoplanin (hamster, 1:500, Novus Biologicals AB15858), DC-LAMP (rat, 1:25, Novus Biologicals, DDX0191P-100), ICAM-1 (rabbit, 1:500, Abcam ab179707), vWF (rabbit 1:1000, Novus Biologicals NB600-586), Endomucin (rat, 1:100, Abcam ab106100), β3-Tubulin (mouse, 1:1000, Abcam ab78078), NeuN (mouse, 1:1000, Novus Biologicals NBP1-92693), GFAP (chicken, 1:1000, Novus Biologicals NBP1-05198), GFP (goat 1:500, Rockland, 600-101-215)

### Western blotting

Animal was perfused with PBS and the tissue of interest was dissected. Tissue (∼5mg) was homogenized in 350ul RIPA buffer supplemented with Protease Inhibitor Cocktail (Roche). Lysates were centrifuged at 12,000 g for 15 min at 4 °C. The supernatant was collected for downstream analysis and flash frozen prior to storage at -80°C. Protein concentration was determined using the Pierce BCA Protein Assay Kit. Protein (30ug) was electrophoresed on a NuPAGE™ 4 to 12% gel and then transferred to a nitrocellulose membrane by the iBlot2 system (Thermo Fisher Scientific) then blocked in 5% nonfat dry milk-TBST for 1 h. Primary antibodies (diluted in 5% milk-TBST) were incubated at 4 °C overnight with gentle agitation, and membranes were then washed three times (5 min each) in TBST. Fluorescent-conjugated secondary antibodies (diluted in 5% milk-TBST) were applied at room temperature for 1 h with gentle agitation, and membranes were then washed five times (5 min each) in TBST. Secondary antibodies were detected using the Licor Imager. Total protein counterstain was used for loading analysis (Revert, Licor 926-11011).

Primary antibodies used for immunoblotting: pan-ACE2 (1:1000; R&D Systems; AF933), hACE2 (1:1000; Atlas Antibodies; AMAB91259), β-Actin (1:1000; Cell Signaling; 4970).

### RNA in situ hybridization

RNAscope *in situ* hybridization (ISH) was performed on decalcified paraffin tissue sections with RNAscope fluorescent multiplex reagent kit (ACD, 320850) according to the manufacturer’s protocol. Briefly, sections were baked at 60 °C for 30 mins, deparaffinized, treated with hydrogen peroxide, followed by antigen retrieval in target retrieval buffer for 15 min, and protease treatment before incubation with RNAscope probe for 2 hours at 40 °C . After probe hybridization, RNA signal was amplified with Amp1, Amp2 and Amp3, followed by HRP-C3 incubation. Color development was performed with TSA plus Cyanine 3 kit (Perkinelmer, NEL744B001KT). The following RNAscope probe was used: nCoV2019-S (ACD, 848561-C3).

For dual ISH and IHC, immunofluorescence staining was carried out after RNAscope ISH according to the manufacturer’s protocol (ACD). Briefly, after ISH, sections were washed with PBS-T (0.1% Tween-20), blocked with 10% normal donkey serum at RT for 1 hour. Sections were then incubated with anti-NFH (chicken 1:5000, Aves Labs, NFH) and anti-KRT8 (rabbit 1:500, Sigma-Aldrich, SAB4501654) overnight at 4°C, followed by incubation with secondary antibodies.

### Antibody validation

ACE2 antibodies were validated for IHC and Western blotting using hACE2^del/y^ tissues. Nucleocapsid and RNAscope detection of SARS-CoV-2 was performed simultaneously on wildtype mouse tissue also infected with virus contemporaneously to experimental animals with tissue sections on the same slide to ensure identical staining conditions.

### Statistics

Animals were inoculated with SARS-CoV-2 in blinded fashion without knowledge of genotypes. At the time of harvest, hACE2 animals could not be handled in a blinded manner given moribund appearance compared to wildtype. All animals were included in analyses except those that spontaneously died prior to harvest where histology could no longer be obtained or animals where the initial viral inoculation was observed to be inconsistent or concern for failure. Unless specifically noted, all animals used in this study were male (ACE2 is X-linked) given the excessive number of animals that would need to be generated to achieve homozygous females with additional Cre-drivers. Foxg1^Cre^; LSL-hACE2^+/0^ experiments in Figure 9 and 10 were done with equal numbers of male and female mice. Prior determination of sample sizes could not be performed given the completely unknown phenotype of our novel genetic models. Sample sizes were pre-determined to be at least three animals per genotype, which was sufficient to achieve statistical significance given the high reproducibility of our models and large effect size. Reproducibility of our findings was rigorously confirmed by infecting our novel genetic models at two separate ABSL-3 facilities with independent experimenters. Randomization of animals was not performed, favoring use of littermates to control for the mixed strain background and given the complexity of the genetic intercrosses. Statistical tests used to determine significance are described in the figure legends. Graph generation and statistical analyses performed with GraphPad Prism 9.2.0. All t-tests performed were two-tailed. All one-way ANOVA tests were performed with Sidak’s correction for multiple comparisons. Survival curve statistics were performed with log-rank Mantel Cox tests.

---

**Extended Data Figure 1.**
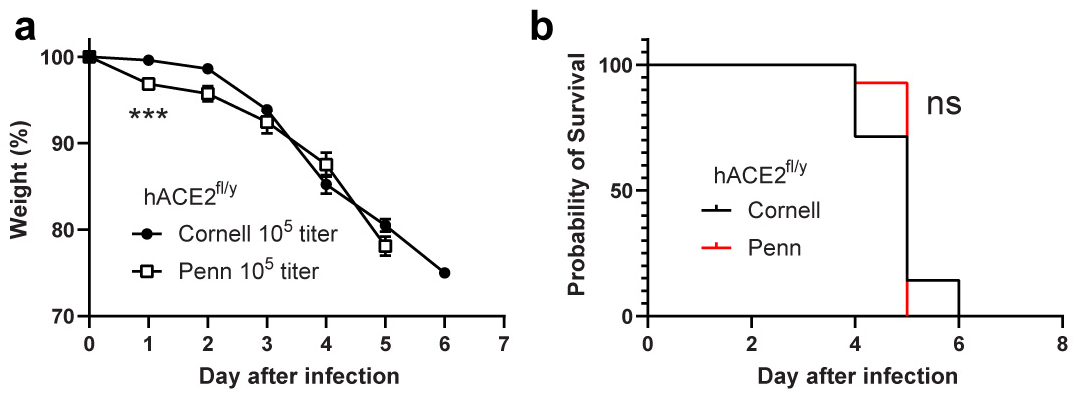
Survival following SARS-CoV-2 infection of *hACE2*^fl/y^ mice in two facilities. **a-b**, Weight loss and survival of *hACE2*^fl/y^ mice were measured after infection with 10^5^ SARS-CoV-2 virus at the Cornell or Penn animal biosafety level 3 (ABSL3) facilities. N=7 (Cornell) and 14 (Penn) from 3 independent experiments. ***p<0.001 by unpaired two-tailed t-test, ns p>0.05 by log-rank Mantel Cox test.

**Extended Data Figure 2.**
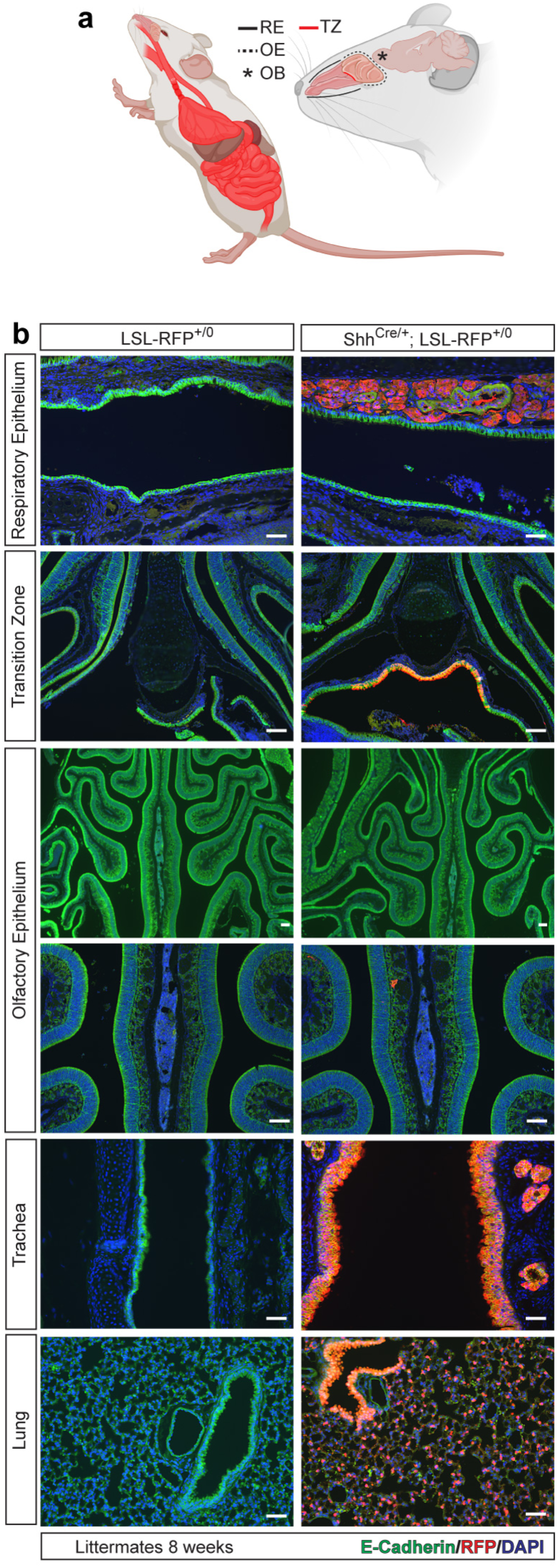
Shh^Cre^ is active in lower respiratory epithelium but spares the nasal cavity epithelium. **a,** Sites of Shh^Cre^ activity in respiratory and gut epithelium are shown in red. RE (respiratory epithelium), OE (olfactory epithelium), OB (olfactory bulb), TZ (transition zone). * indicates olfactory bulb. Lines trace regions of nasal cavity epithelium. **b,** Lineage trace of Shh^Cre^ activity in upper and lower respiratory epithelium using a Cre-activated, tdTomato allele (LSL-RFP). Immunohistochemistry using E-cadherin (epithelium) and RFP (Cre reporter) antibodies is shown. N=3 per genotype. Scale bars 100 µm.

**Extended Data Figure 3.**
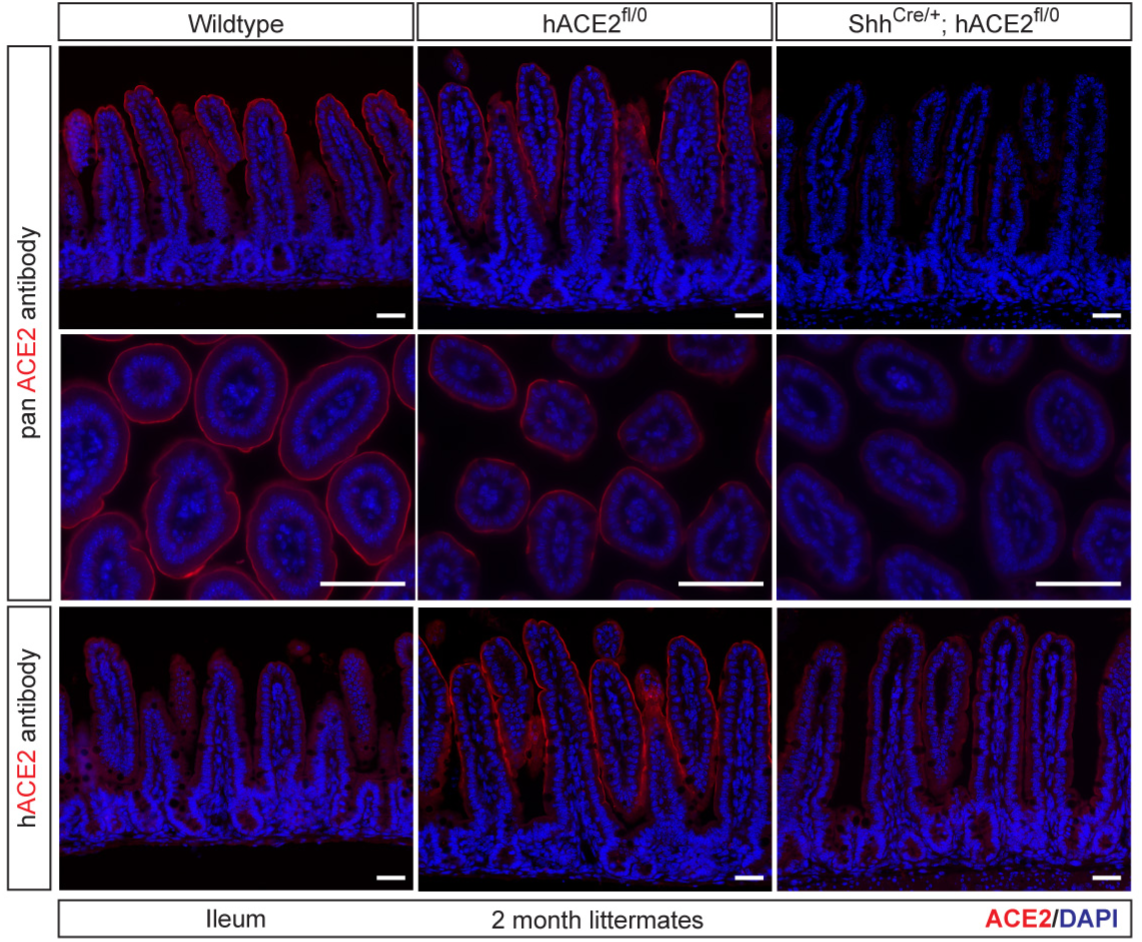
Reduced small intestine epithelial expression of hACE2 in Shh^Cre^; *hACE2*^fl/y^ mice. Immunohistochemistry of wild type, *hACE2*^fl/y^ and Shh^Cre^; *hACE2*^fl/y^ small intestine using antibodies that recognize both human and mouse ACE2 (pan-ACE2) and only human ACE2 (hACE2) are shown. Scale bars 50 µm.

**Extended Data Figure 4.**
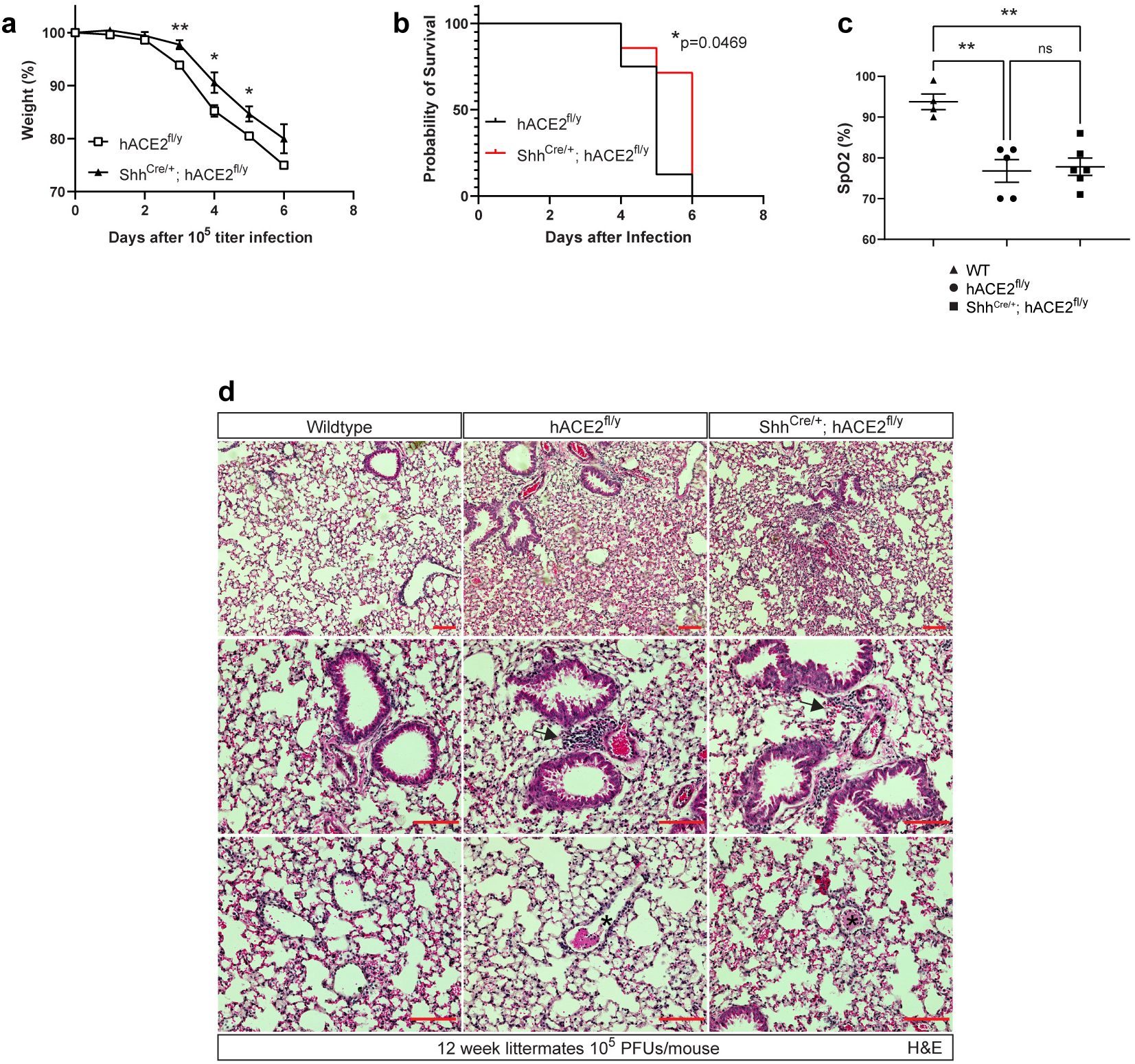
Acute lung injury, hypoxemia and lethality in *hACE2*^fl/y^ and Shh^Cre/+^; *hACE2*^fl/y^ mice after infection with SARS-CoV-2. **a-b**, Weight loss and survival of *hACE2*^fl/y^ and Shh^Cre/+^; *hACE2*^fl/y^ mice after infection with 10^5^ PFU of SARS-CoV-2. N=8 (*hACE2*^fl/y^) and 7 (Shh^Cre/+^; *hACE2*^fl/y^), two independent experiments. Note: data for *hACE2*^fl/y^ are the same as those shown in Figure 2d, e because Shh^Cre/+^; *hACE2*^fl/y^ animals were littermates of those animals. **c**, Pulse oximetry measured in wild-type (WT), *hACE2*^fl/y^ and Shh^Cre/+^; *hACE2*^fl/y^ mice at time of harvest 5-6 days after exposure to 10^5^ PFU of SARS-CoV-2 virus. **d**, Hematoxylin-eosin (H&E) staining of wild-type (WT), *hACE2*^fl/y^ and Shh^Cre/+^; *hACE2*^fl/y^ lung tissue 6 days after exposure to 10^5^ PFU of SARS-CoV-2 virus. Arrows, sites of broncho-vascular immune cell infiltrate. Asterisks, intravascular thrombus Representative of N=3 animals per genotype. Scale bars 100 µm. *p<0.05, **p<0.01, ns p>0.05 by unpaired two-tailed t-test or log-rank Mantel Cox test.

**Extended Data Figure 5.**
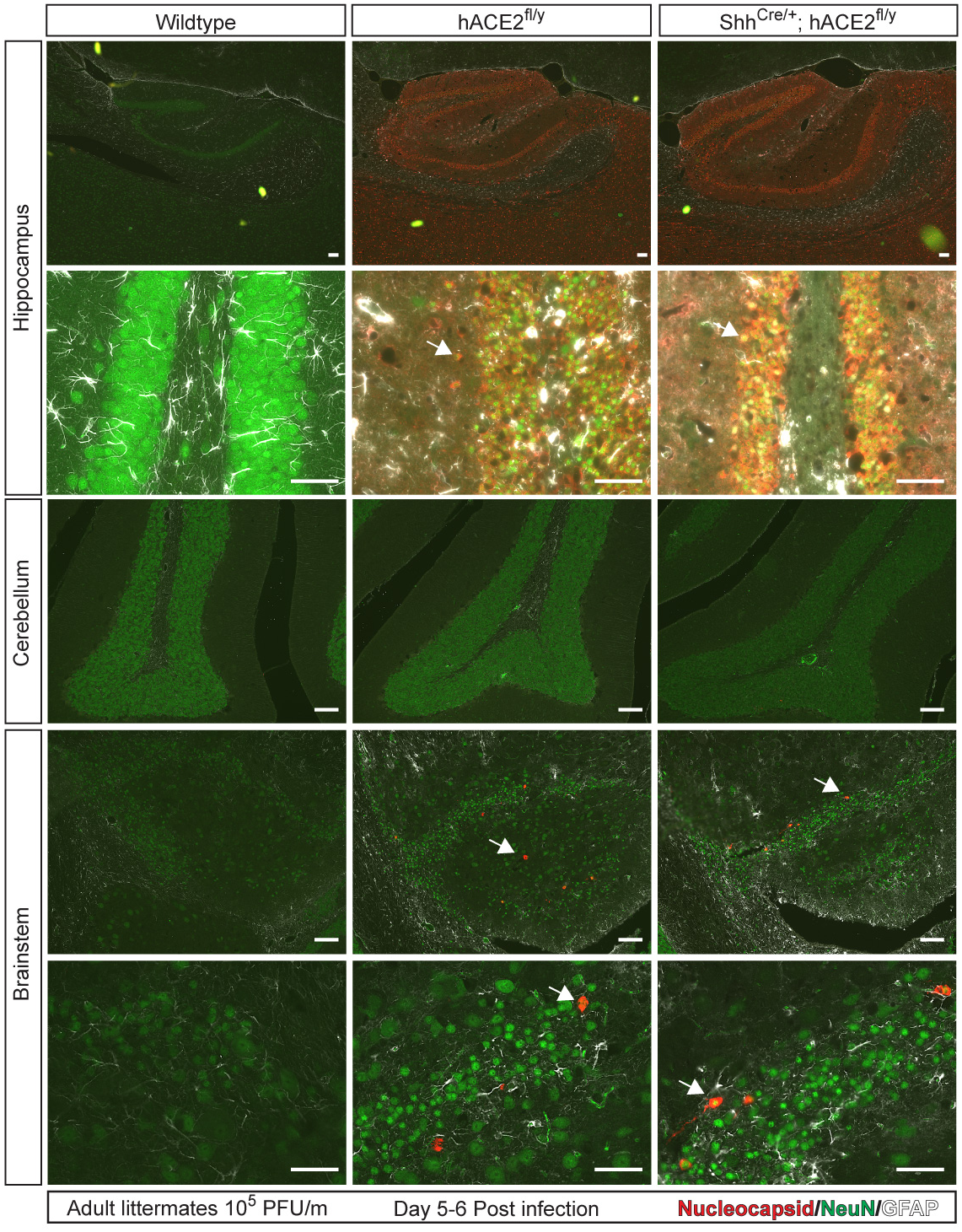
Extensive SARS-CoV-2 detection in hippocampus but minimal detection in neurons of the cerebellum and brainstem. Immunohistochemistry of wildtype, *hACE2*^fl/y^ and Shh^Cre/+^; *hACE2*^fl/y^ hippocampus, cerebellum and brainstem with antibody staining for viral nucleocapsid, neurons (NeuN), and glial cells (GFAP). Arrows indicate NeuN positive neurons with colocalized nucleocapsid staining. N=3 per genotype. Scale bars 100 µm (top three rows), 50 µm (bottom row).

**Extended Data Figure 6.**
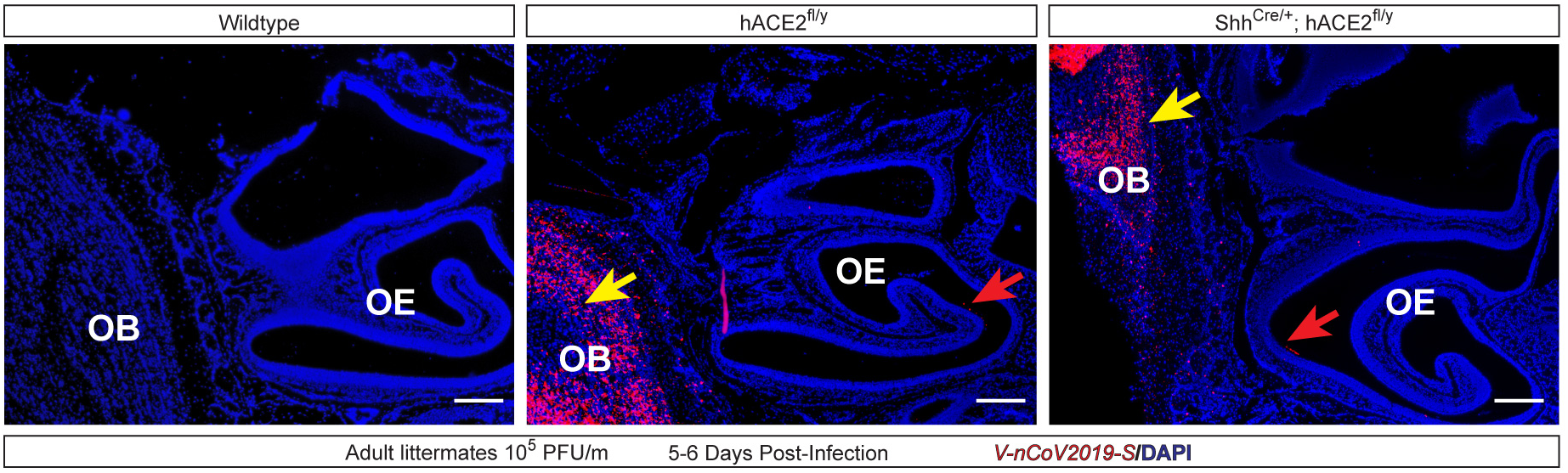
SARS-CoV-2 infection of the olfactory epithelium and olfactory bulb detected by *in situ* hybridization. *In situ* hybridization of SARS-CoV-2 mRNA in wildtype, *hACE2*^fl/y^ and Shh^Cre/+^; *hACE2*^fl/y^ mice infected with 10^5^ viral titer was performed on tissue samples harvested 5-6 days post infection. Yellow arrows indicate positive hybridization signal in the olfactory bulb (OB), Red arrows indicate positive hybridization signal in the olfactory epithelium (OE). N=3 per genotype. Scale bars 250 µm.

**Extended Data Figure 7.**
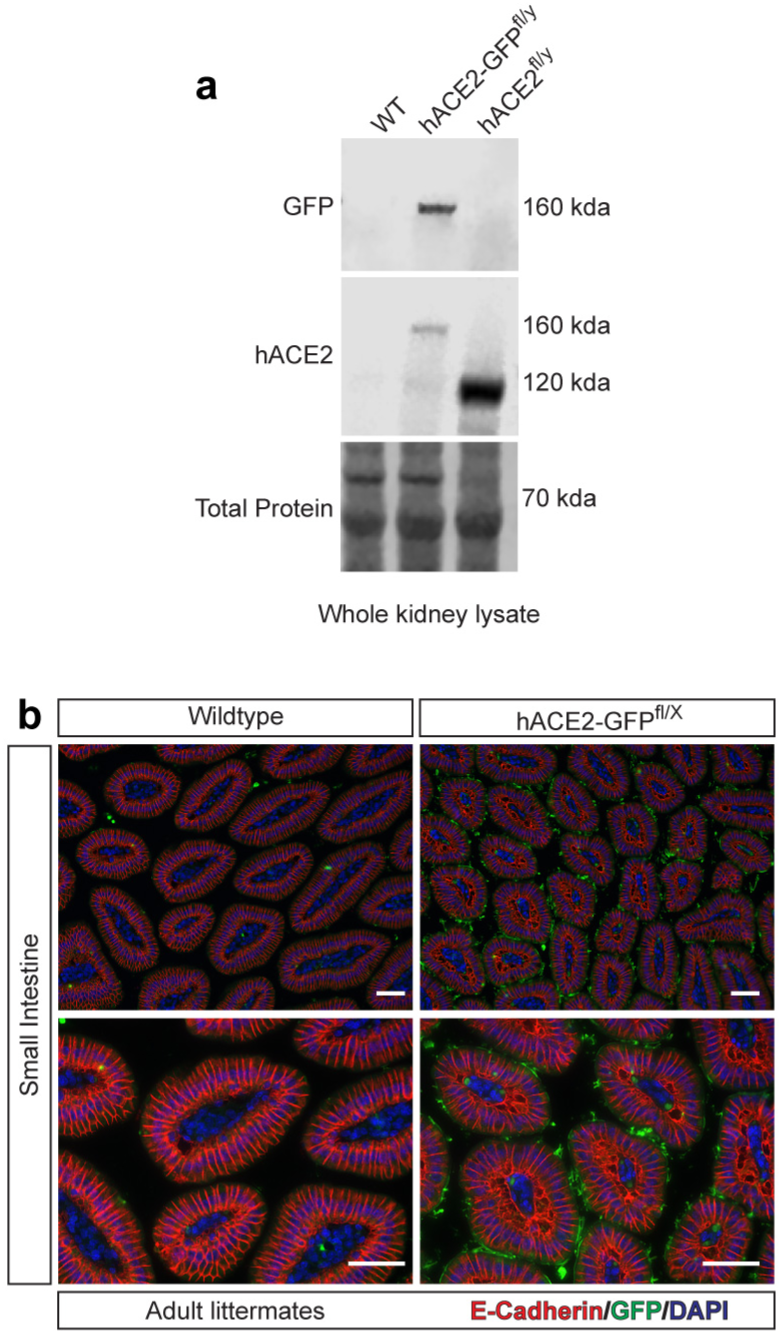
Expression of hACE2-GFP protein in *hACE2-GFP*^fl/y^ kidney and small intestine. **a**, Immunoblotting of whole kidney lysate from wildtype, *hACE2-GFP*^fl/y^, and *hACE2*^fl/y^ animals using anti-GFP antibodies and antibodies specific for human-ACE2 (hACE2). Revert^TM^ total protein counterstain used for loading control. N=3 per genotype, two independent experiments. **b,** Immunohistochemistry of small intestine from wildtype and *hACE2-GFP*^fl/X^ animals using antibodies that recognize GFP and E-cadherin (epithelium). N=3 per genotype, three independent experiments. Scale bars 50 µm.

**Extended Data Figure 8.**
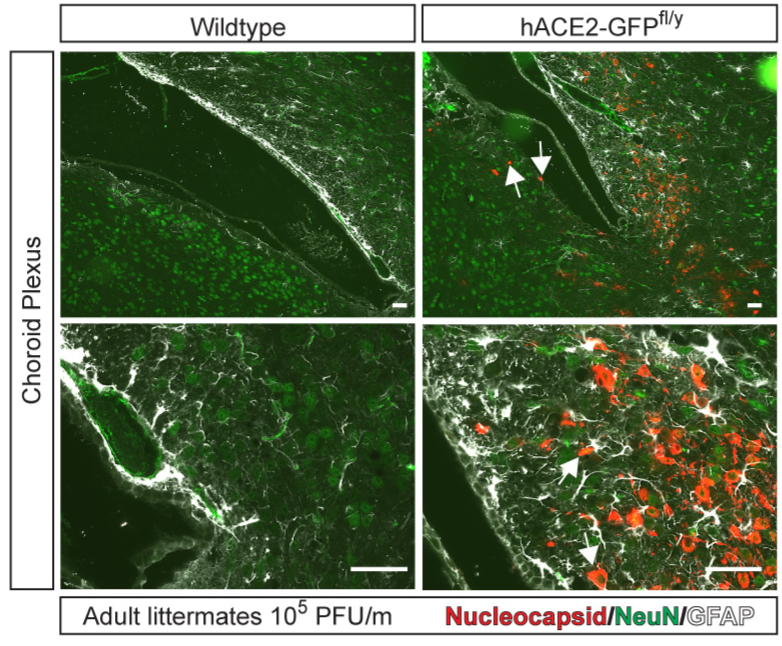
SARS-CoV-2 detection in neurons adjacent to the choroid plexus of hACE2-GFP^fl/y^ mice. Immunohistochemistry of the choroid plexus along the lateral ventricle of wildtype and hACE2-GFP^fl/y^ mice 6 days post-infection with 10^5^ viral titer per mouse. Staining for viral nucleocapsid, neuronal NeuN, and glial GFAP. Arrows indicate nucleocapsid staining colocalized with NeuN. Red dotted lines trace ependymal cells of the choroid plexus. Results representative of N=3 per genotype. Scale bars 50 µm.

**Extended Data Figure 9.**
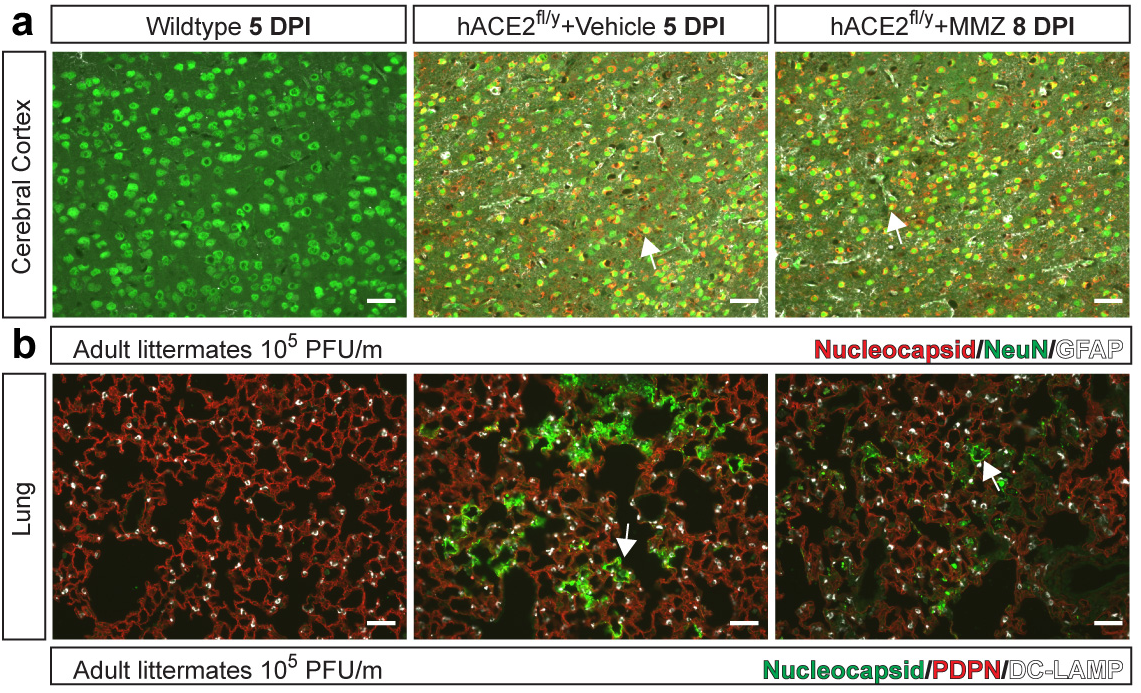
Failure of methimazole to prevent lethal SARS-CoV-2 infection is associated with brain infection. **a-b,** Cerebral cortex (a) and lung (b) immunohistochemistry of *hACE2*^fl/y^ animals treated with vehicle or methimazole (MMZ) followed by infection with 10^5^ viral titer. The tissues shown on the far right (“*hACE2*^fl^ + MMZ 8 DPI”) are from a *hACE2*^fl^ mouse that exhibited weight loss and required euthanasia 8 days following infection with 10^5^ SARS-CoV-2 virus despite pre-treatment with MMZ. Cerebral cortex stained with viral nucleocapsid, neuronal NeuN and glial cell GFAP. Lung stained with viral nucleocapsid, alveolar type I cells PDPN, and alveolar type II cell DC-LAMP. Arrows indicate nucleocapsid staining colocalized with NeuN+ neurons or PDPN+ AT1 cells. N=4 per genotype except for N=1 (MMZ treated). Scale bars 50 µm.

**Extended Data Figure 10.**
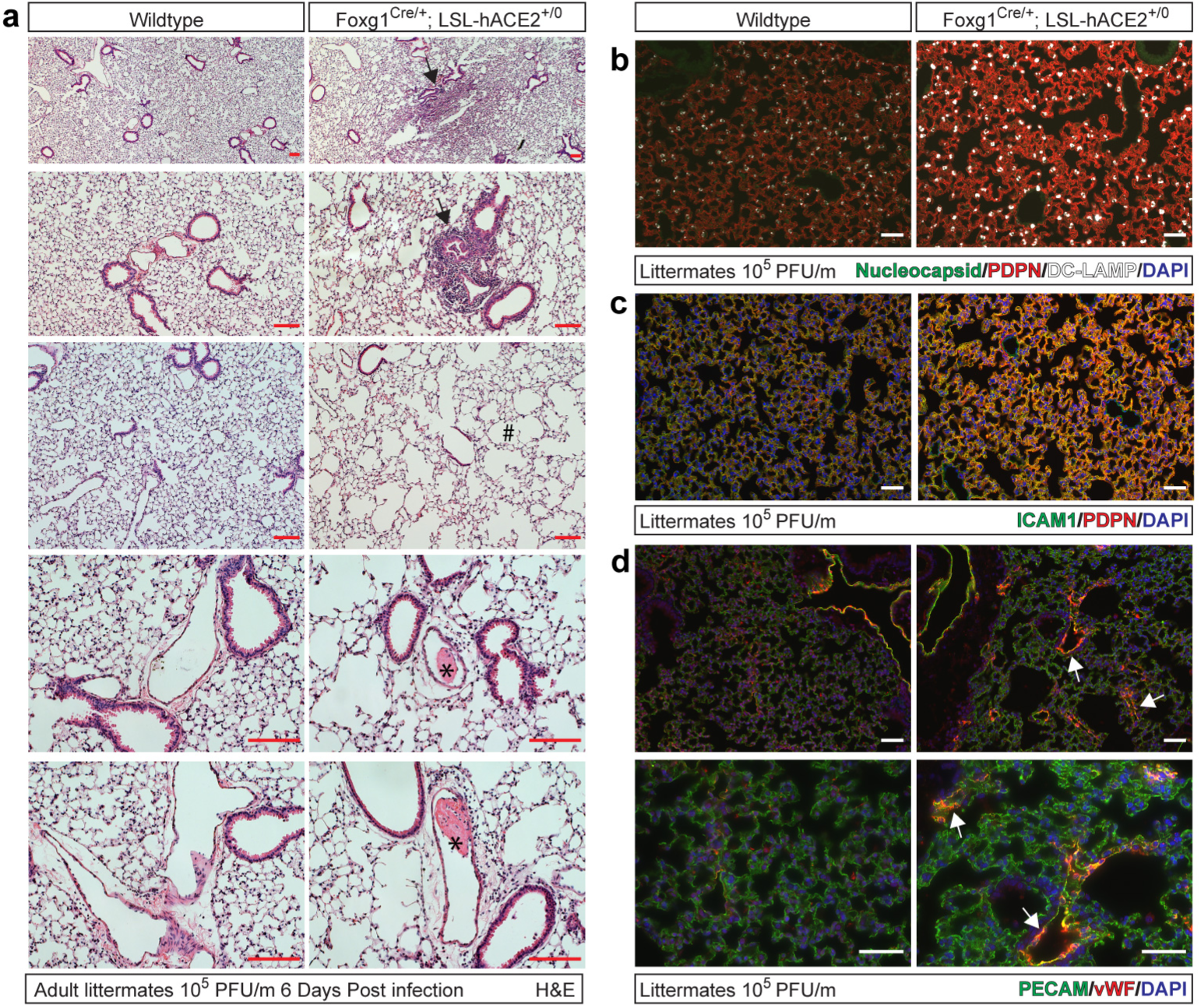
SARS-CoV-2 infection of OE and neurons is sufficient to confer pulmonary inflammation and intravascular thrombosis. **a**, Hematoxylin-eosin (H&E) staining of wild-type and Foxg1^Cre/+^; LSL-hACE2^+/0^ lung tissue 6 days after exposure to 10^5^ PFU of SARS-CoV-2 virus. Arrows, sites of focal consolidation. Asterisks, intravascular thrombi. Hashtag, acute emphysematous changes. Representative of N=4 animals per genotype. Scale bars 100 µm. **b,** Immunohistochemistry of wild-type and Foxg1^Cre/+^; LSL-hACE2^+/0^ lung tissue 6 days after exposure to 10^5^ PFU of SARS-CoV-2 virus using antibodies against viral nucleocapsid, alveolar type 1 PDPN, and alveolar type II DC-LAMP. Representative of N=4 animals per genotype, Scale bars 50 µm. **c-d**, Immunohistochemistry of wild-type and Foxg1^Cre/+^; LSL-hACE2^+/0^ lung tissue 6 days after exposure to 10^5^ PFU of SARS-CoV-2 virus using antibodies against Intracellular adhesion marker-1 (ICAM-1) and Podoplanin (PDPN), or von Willebrand’s Factor (vWF) and PECAM (endothelial cells). Arrowheads in d identify vWF-positive microvasculature of the lung in Foxg1^Cre/+^; LSL-hACE2^+/0^ animals. Representative of N=3-4 animals per genotype. Scale bars, 50 µm.

## Notes

### Competing Interest Statement

The authors have declared no competing interest.

